# Deep protein methylation profiling by combined chemical and immunoaffinity approaches reveals novel PRMT1 targets

**DOI:** 10.1101/538686

**Authors:** Nicolas G. Hartel, Brandon Chew, Jian Qin, Jian Xu, Nicholas A. Graham

## Abstract

Protein methylation has been implicated in many important biological contexts including signaling, metabolism, and transcriptional control. Despite the importance of this post-translational modification, the global analysis of protein methylation by mass spectrometry-based proteomics has not been extensively studied due to the lack of robust, well-characterized techniques for methyl peptide enrichment. Here, to better investigate protein methylation, we compared two methods for methyl peptide enrichment: immunoaffinity purification (IAP) and high pH strong cation exchange (SCX). Using both methods, we identified 1,720 methylation sites on 778 proteins. Comparison of these methods revealed that they are largely orthogonal, suggesting that the usage of both techniques is required to provide a global view of protein methylation. Using both IAP and SCX, we then investigated changes in protein methylation downstream of protein arginine methyltransferase 1 (PRMT1). PRMT1 knockdown resulted in significant changes to 127 arginine methylation sites on 78 proteins. In contrast, only a single lysine methylation site was significantly changed upon PRMT1 knockdown. In PRMT1 knockdown cells, we found 114 MMA sites that were either significantly downregulated or upregulated on proteins enriched for mRNA metabolic processes. PRMT1 knockdown also induced significant changes in both asymmetric dimethyl arginine (ADMA) and symmetric dimethyl arginine (SDMA). Using characteristic neutral loss fragmentation ions, we annotated dimethylarginines as either ADMA or SDMA. Through integrative analysis of methyl forms, we identified 18 high confidence PRMT1 substrates and 12 methylation sites that are scavenged by other non-PRMT1 arginine methyltransferases in the absence of PRMT1 activity. We also identified one methylation site, HNRNPA1 R206, which switched from ADMA to SDMA upon PRMT1 knockdown. Taken together, our results suggest that deep protein methylation profiling by mass spectrometry requires orthogonal enrichment techniques to identify novel PRMT1 methylation targets and highlight the dynamic interplay between methyltransferases in mammalian cells.

## INTRODUCTION

Protein post-translational modifications (PTMs) regulate diverse biological processes and provide additional complexity to proteins beyond their initial primary sequence (1). Protein methylation was first identified over 50 years ago on both arginine (2) and lysine (3) residues, yet these PTMs are among the least studied compared to other modifications such as phosphorylation, acetylation, and ubiquitination (4). Regardless, recent studies have identified protein arginine methylation as an important regulator of signal transduction (5–7), metabolism (8, 9), cell cycle (10), and transcriptional control (11–13).

The mammalian genome encodes nine protein arginine methyltransferases (PRMTs) and approximately 50 lysine methyltransferases (KMTs). Both PRMTs and KMTs use S-adenosylmethionine (SAM) as a methyl donor to methylate either the guanidino nitrogens of arginine or the ε-amino group of lysine. The complexity of protein methylation is enhanced by the fact that both methyl-arginine and methyl-lysine occur in three distinct forms. Arginine exists in monomethyl (MMA), asymmetric dimethyl (ADMA), or symmetric dimethyl (SDMA) forms, whereas lysine exists in monomethyl (Kme1), dimethyl (Kme2), or trimethyl (Kme3) forms. PRMTs can be divided into two categories based on which type of arginine methylation they catalyze: Type I PRMTs catalyze MMA and ADMA (PRMT1, PRMT3, PRMT4, PRMT6, and PRMT8) (14), whereas Type II catalyze MMA and SDMA (PRMT5 and PRMT9) and Type III catalyze MMA only (PRMT7) (15).

One reason why the study of protein methylation has lagged behind other PTMs is a lack of robust methyl peptide enrichment strategies (16). Compared to strategies for enrichment of phospho-peptides with TiO_2_ (17) or IMAC (18) or for enrichment of glycosylated peptides with hydrophilic interaction chromatography (HILIC) (19), techniques for enriching methyl peptides have not been as widely adopted. Enrichment of methyl peptides is more difficult than many other PTMs because methylation does not add significant steric bulk or change the charge of the amino acids (20). Despite this difficulty, advances in methyl peptide enrichment for mass spectrometry have been made using immunoaffinity enrichment (IAP) with antibodies that recognize various forms of protein methylation (21–26). By combining IAP against MMA with sample fractionation, as many as 8,000 MMA sites have been identified in human cells (24). Other enrichment strategies for methyl peptides include high pH strong cation exchange (SCX), chemical labeling, HILIC, and engineered MBT domains that bind methylated proteins (27–31). High pH SCX relies on missed cleavage by trypsin of methylated arginine and lysine residues which results in methyl peptides with higher positive charge that can be enriched by SCX. Together, these enrichment techniques have begun to shed insight into the global regulation of protein methylation, but there has been no extensive comparison of enrichment methods to present a global picture of protein methylation.

Here, to better study protein methylation, we compared two methyl peptide enrichment strategies, high pH SCX and IAP. Notably, comparison of high pH SCX and IAP revealed that these methods are largely orthogonal and quantitatively reproducible, suggesting that both methods are required for global analysis of protein methylation. We then used both methyl proteomics methods in parallel to investigate the PRMT1 methylome. Knockdown of PRMT1 with shRNA led to significant changes in both MMA and DMA sites, primarily on RNA binding proteins. Additionally, examination of MS/MS spectra confirmed that ADMA and SDMA peptides can be distinguished by neutral ion loss from methylarginine (16, 32–34). Through integrative analysis of MMA and DMA, we identified a list of 18 PRMT1 substrates and 12 substrates scavenged by other PRMTs in the absence of PRMT1 activity. Taken together, our results describe a general method for deep profiling of protein methylation and identify novel potential MMA and ADMA methylation targets of PRMT1.

## EXPERIMENTAL PROCEDURES

### Cell Culture

LN229 cells and HEK 293T cells expressing short hairpin RNA (shRNA) against PRMT1 or control were grown in DMEM media (Corning) supplemented with 10% FBS (Omega Scientific) and 100 U/mL penicillin/streptomycin (Thermo Scientific). Cells were cultured at 37 °C in humidified 5% CO_2_ atmosphere. Generation of HEK 293T cells stably expressing shRNA against PRMT1 or control were previously described (5). shPRMT1 and shControl cells were cultured with 4 ug/mL puromycin to maintain selection.

### Cell Lysate Preparation

Cells were washed with PBS, scraped, and lysed in 50 mM Tris pH 7.5, 8M urea, 1 mM activated sodium vanadate, 2.5 mM sodium pyrophosphate, 1 mM β-glycerophosphate, and 100 mM sodium phosphate. Protein concentrations were measured by bicinchoninic assay. Lysates were sonicated and cleared by high speed centrifugation and then filtered through 0.22 um filter. Proteins were reduced, alkylated, and quenched with 5 mM dithiothreitol, 25 mM iodoacetamide, 10 mM dithiothreitol, respectively. Lysates were four-fold diluted in 100 mM Tris pH 8.0 and digested with trypsin at a 1:100 ratio and then quenched with addition of trifluoroacetic acid to pH 2. Peptides were purified using reverse-phase Sep-Pak C18 cartridges (Waters) and eluted with 30% acetonitrile, 0.1% TFA and then dried by vacuum. Dried peptides were subjected to high pH strong cation exchange or antibody immunoaffinity purification.

### Immunoblot Analysis

Cells were lysed in modified RIPA buffer (50 mM Tris–HCl (pH 7.5), 150 NaCl, 50 mM β-glycerophosphate, 0.5 mM NP-40, 0.25% sodium deoxycholate, 10 mM sodium pyrophosphate, 30 mM sodium fluoride, 2 mM EDTA, 1 mM activated sodium vanadate, 20 μg/ml aprotinin, 10 μg/ml leupeptin, 1 mM DTT, and 1 mM phenylmethylsulfonyl fluoride). Whole-cell lysates were resolved by SDS–PAGE on 4–15% gradient gels and blotted onto nitrocellulose membranes (Bio-Rad). Membranes were blocked for 1 h in nonfat-milk, and then incubated with primary and secondary antibodies overnight and for 2 h, respectively. Blots were imaged using the Odyssey Infrared Imaging System (Li-Cor). Primary antibodies used for Western blot analysis were: mono-methyl arginine (8015, Cell Signaling), asymmetric di-methyl arginine motif (13522, Cell Signaling), symmetric di-methyl arginine motif (13222, Cell Signaling), PRMT1 (2449, Cell Signaling), and anti-β-actin (10081-976, Proteintech).

### High pH Strong Cation Exchange (SCX)

As described previously (28), in brief, 1 mg of digested protein was resuspended in loading buffer (60% acetonitrile, 40% BRUB (5 mM phosphoric acid, 5 mM boric acid, 5 mM acetic acid, pH 2.5) and incubated with high pH SCX beads (Sepax) for 30 minutes, washed with washing buffer (80% acetonitrile, 20% BRUB, pH 9), and eluted into five fractions using elution buffer 1 (60% acetonitrile, 40% BRUB, pH 9), elution buffer 2 (60% acetonitrile, 40% BRUB, pH 10), elution buffer 3 (60% acetonitrile, 40% BRUB, pH 11), elution buffer 4 (30% acetonitrile, 70% BRUB, pH 12), and elution buffer 5 (100% BRUB, 1M NaCl, pH 12). Eluates were dried, resuspended in 1% trifluoroacetic acid and desalted on STAGE tips (35) with 2 mg of HLB material (Waters) loaded onto 300 uL tip with a C8 plug (Empore, Sigma). *Immunoaffinity Purification (IAP)* — 10 mg of digested proteins were dissolved in 1X immunoprecipitation buffer (50 mM MOPS, 10 mM Na_2_HPO_4_, 50 mM NaCl, pH 7.2, Cell Signaling). Modified symmetric dimethyl arginine peptides, asymmetric dimethyl arginine peptides, and monomethyl arginine peptides were immunoprecipitated by addition of 40 uL of PTMScan Symmetric Di-Methyl Arginine Motif Kit (13563, Cell Signaling), PTMScan Asymmetric Di-Methyl Arginine Motif Kit (13474, Cell Signaling), and PTMScan Mono-Methyl Arginine Motif Kit (12235, Cell Signaling), respectively. Modified methyl lysine peptides were enriched with PTMScan Pan-Methyl Lysine Kit (14809). Lysates were incubated with PTMScan motif kits for 2 hours at 4 °C on a rotator. Beads were centrifuged and washed two times in 1X immunoprecipitation buffer followed by three washes in water, and modified peptides were eluted with 2 x 50 uL of 0.15% TFA and desalted on STAGE tips with C18 cores (Empore, Sigma). Enriched peptides were resuspended in 50 mM ammonium bicarbonate (Sigma) and subjected to a second digestion with trypsin for 2 hours per the manufacturer’s recommendation, acidified with trifluoroacetic acid to pH 2 and desalted on STAGE tips.

### Mass Spectrometric Analysis

All LC-MS experiments were performed on a nanoscale UHPLC system (EASY-nLC1200, Thermo Scientific) connected to an Q Exactive Plus hybrid quadrupole-Orbitrap mass spectrometer equipped with a nanoelectrospray source (Thermo Scientific). Peptides were separated by a reversed-phase analytical column (PepMap RSLC C18, 2 μm, 100 Å, 75 μm X 25 cm) (Thermo Scientific). For high pH SCX fractions a “Short” gradient was used where flow rate was set to 300 nl/min at a gradient starting with 0% buffer B (0.1% FA, 80% acetonitrile) to 29% B in 142 minutes, then washed by 90% B in 10 minutes, and held at 90% B for 3. The maximum pressure was set to 1,180 bar and column temperature was constant at 50 °C. For IAP samples a “Slow” gradient was used where flow rate was set to 300 nl/min at a gradient starting with 0% buffer B to 25% B in 132 minutes, then washed by 90% B in 10 minutes. Dried SCX fractions were resuspended in buffer A and injected as follows, E1: 1.5 μL/60 μL, E2-5: 5 μL/6 μL. IAP samples were resuspended in 7 μL and 6.5 μL was injected. The effluent from the HPLC was directly electrosprayed into the mass spectrometer. Peptides separated by the column were ionized at 2.0 kV in the positive ion mode. MS1 survey scans for DDA were acquired at resolution of 70k from 350 to 1,800 m/z, with maximum injection time of 100 ms and AGC target of 1e6. MS/MS fragmentation of the 10 most abundant ions were analyzed at a resolution of 17.5k, AGC target 5e4, maximum injection time 120 ms for IAP samples, 240 ms for SCX samples, and normalized collision energy 26. Dynamic exclusion was set to 30 s and ions with charge 1 and >6 were excluded. The mass spectrometry proteomics data have been deposited to the ProteomeXchange Consortium (http://proteomecentral.proteomexchange.org) via the PRIDE partner repository with the dataset identifier PXD012357 (Username, password: 5WbkZaM5) (36).

### Identification and Quantitation of Peptides

MS/MS fragmentation spectra were searched with Proteome Discoverer SEQUEST (version 2.2, Thermo Scientific) against the in-silico tryptic digested Uniprot *H. sapiens* database with all reviewed with isoforms (release Jun 2017, 42,140 entries). The maximum missed cleavage rate was set to 5 (28). Trypsin was set to cleave at R and K. Dynamic modifications were set to include mono-methylation of arginine or lysine (R/K, +14.01565), di-methylation of arginine or lysine (R/K, +28.0313), tri-methylation of lysine (K, +42.04695), oxidation on methionine (M, +15.995 Da, and acetylation on protein N-terminus (+42.011 Da). Fixed modification was set to carbamidomethylation on cysteine residues (C, +57.021 Da). The maximum parental mass error was set to 10 ppm and the MS/MS mass tolerance was set to 0.02 Da. Peptides with sequence of six to fifty amino acids were considered. Methylation site localization was determined by ptm-RS node in Proteome Discoverer, and only sites with localization probability greater or equal to 75% were considered. The False Discovery Rate threshold was set strictly to 0.01 using Percolator node validated by q-value. Relative abundances of parental peptides were calculated by integration of area-under-the-curve of the MS1 peaks using Minora LFQ node in Proteome Discoverer 2.2. The Proteome Discoverer export peptide groups abundance values were log_2_ transformed, normalized to the corresponding samples median values, and significance was determined using a permutation-based FDR approach in the Perseus environment (37) (release 1.6.2.3) with a q-value FDR of 0.05 and S_0_ value of 0.5.

### Methyl False Discovery Estimation

The “Decoy PSMs” export from Proteome Discoverer 2.2 was filtered for decoy methyl PSMs and the decoy q-values from the Percolator node were extracted and compared to the target methyl PSM q-values. Target methyl PSMs were removed until a 1% FDR was achieved as described (38).

### Neutral Loss Identification in MaxQuant

The modifications SDMA and ADMA were added to MaxQuant’s library with the added mass of dimethyl on arginine and the corresponding neutral loss masses of 31.042 for SDMA and 45.058 for ADMA assigned in the “Neutral Loss” table in Configuration (26). The missed cleavage rate was set to 5 and all other settings were kept unchanged. All RAW files were searched with monomethyl(K/R), ADMA, SDMA, and oxidation of methionine as variable modifications. Carbamidomethylation was kept as a fixed modification. Neutral losses and their masses were extracted from the msms.txt file. Only target methyl peptides that passed the 1% Methyl FDR filter were considered for analysis. An Andromeda cutoff score of 56 was also used to filter spectra to reduce the number of incorrect assignments. A custom R script was used to remove neutral losses that did not have the corresponding b/y ion present (e.g., if y6* but not y6 was present, the neutral loss was removed). A few spectra were confirmed by manual inspection to ensure the accuracy of the Andromeda search. For identified ADMA / SDMA neutral losses, the Andromeda output was matched to Proteome Discoverer data by MS2 scan number.

### Motif Analysis

Motifs were analyzed by MotifX (39) and MOMO from MEME suite (40) to detect statistically significant patterns in methylation sequence data. Two sample motif analysis was performed using Two Sample Logo (41).

## EXPERIMENTAL DESIGN AND STATISTICAL RATIONALE

### Samples

Three LN229 samples were analyzed, two were enriched by high pH SCX enrichment and 1 enriched by mono-methyl arginine immunoaffinity purification. Two SCX samples were each run on the “Long” and “Short” SCX gradients, and the “Short” gradient SCX samples were compared to the single mono-methyl arginine IAP sample. The single mono-methyl arginine IAP sample was injected in two equal amounts on a “Standard” and “Slow” gradient. Four shPRMT1 293T samples and four shControl 293T samples were analyzed and compared. Two of the four 293T samples were enriched by high pH SCX and another two were enriched by sequential immunoaffinity purification incubated with SDMA, ADMA, MMA, and Pan-K PTMScan Kits sequentially. 50ug of whole cell lysates from each sample were collected for immunoblots before SCX or IAP enrichment.

### Replicates

293T cells had two biological replicates each for high pH SCX enrichment and each sequential IAP enrichment.

### Controls and Randomization

293T cells expressing shRNA against PRMT1 were controlled by using 293T expressing scrambled non-targeting short hairpin control RNA to account for biases introduced by stable transfection. LN229 and 293T samples were also randomized by sample prior to LC-MS injection.

### Rationale

Using only two replicates for 293T shPRMT1 and 293T shControl is justified because of a low coefficient of variance for quantified peptides (85% of MMA IAP peptides below 60% CV for shControl and 78% of MMA IAP peptides below 60% CV for shPRMT1). A third of each dataset is also below 20% CV which indicates small variability between biological replicates. Two replicates for the 293T SCX replicates is also justified due to fractions E1, E2, and E3 having median CV’s of 21, 26, and 54 respectively for shControl samples and 32, 66, and 70 for the shPRMT1 samples. Fractions E1,2, and 3 contributed to 95% of quantified methyl peptides for high pH SCX. Sample abundances from Minora Label Free Quantitation through Proteome Discoverer 2.2 were log_2_ transformed, median normalized by sample, then subject to a permutation-based FDR approach in Perseus software with a q-value FDR of 0.05 and S_0_ of 0.5. All datasets showed a normal distribution of abundances after log_2_ transformation. Statistical significance was given to differential peptide pairs with q-value < 0.05 and required confident methyl site localization of 75% or more through ptm-RS node on Proteome Discoverer. Missing values across samples were excluded and only peptides confidently quantified in all replicates were considered for the permutation-based FDR calculation. For SCX methyl peptides identified in multiple fractions, the largest raw abundance was used to determine which fraction to use to calculate peptide abundance. Taken together, the low variability of biological replicates and rigorous statistical thresholds (methyl peptide discovery FDR = 1% and permutation FDR = 5%) allow confident differential analysis of methyl peptides.

## RESULTS

### SCX and IAP enrich methyl peptides and target different subsets of protein methylome

We first optimized the LC gradients for high pH SCX and IAP to enhance the detection of hydrophilic methylated peptides. By shortening and lengthening the gradients for SCX and IAP, respectively, modest improvements were made in instrument time and number of unique methyl peptides identified (Fig. S1-2, Tables S1-2). Next, we applied our workflow to lysates from 293T cells expressing short hairpin RNA against PRMT1 (shPRMT1) or a negative control (shControl) (Fig. 1A). Across all experiments, we identified 1,720 methylation sites on 778 proteins with a strict 1% methyl-peptide FDR (Tables S3-S7). A summary of the peptide spectral matches (PSMs) from each technique is provided in Table 1. Each technique enriched the expected type of arginine methylation: MMA IAP identified primarily MMA peptides, ADMA IAP and SDMA IAP identified primarily dimethyl arginine (DMA) peptides, and SCX identified primarily DMA peptides (Fig. 1B). PanK and SCX both identified an even distribution of monomethyl lysine (Kme1), dimethyl lysine (Kme2), and trimethyl lysine (Kme3) PSMs. Similar to previous reports, we identified roughly 5 times as many methyl-arginine sites as methyl-lysine sites (42). Notably, the overlap of unique methyl peptides enriched by SCX and IAP was relatively low for all types of arginine and lysine methylation (Fig. 1C), demonstrating that these enrichment techniques target different methyl arginine peptides. Gene ontology analysis of methyl peptides identified by SCX and IAP demonstrated that both techniques were highly enriched for RNA binding proteins (Fig. 1D), in agreement with known properties of methyl proteins (11, 24, 43–47). In addition, IAP enriched proteins related to DNA binding and transcription factor binding, whereas SCX enriched proteins related to nucleoside-triphosphatase activity and hydrolase activity. Comparison of the number of methyl arginine sites per PSM revealed that IAP generally enriched singly methylated peptides while SCX enriched multi-methylated peptides, with some SCX PSMs containing up to four methylation sites (Fig. 1E). Taken together, this data suggests that the usage of both IAP and SCX methods is required to achieve a more complete coverage of the protein arginine methylome.

**Figure 1:**
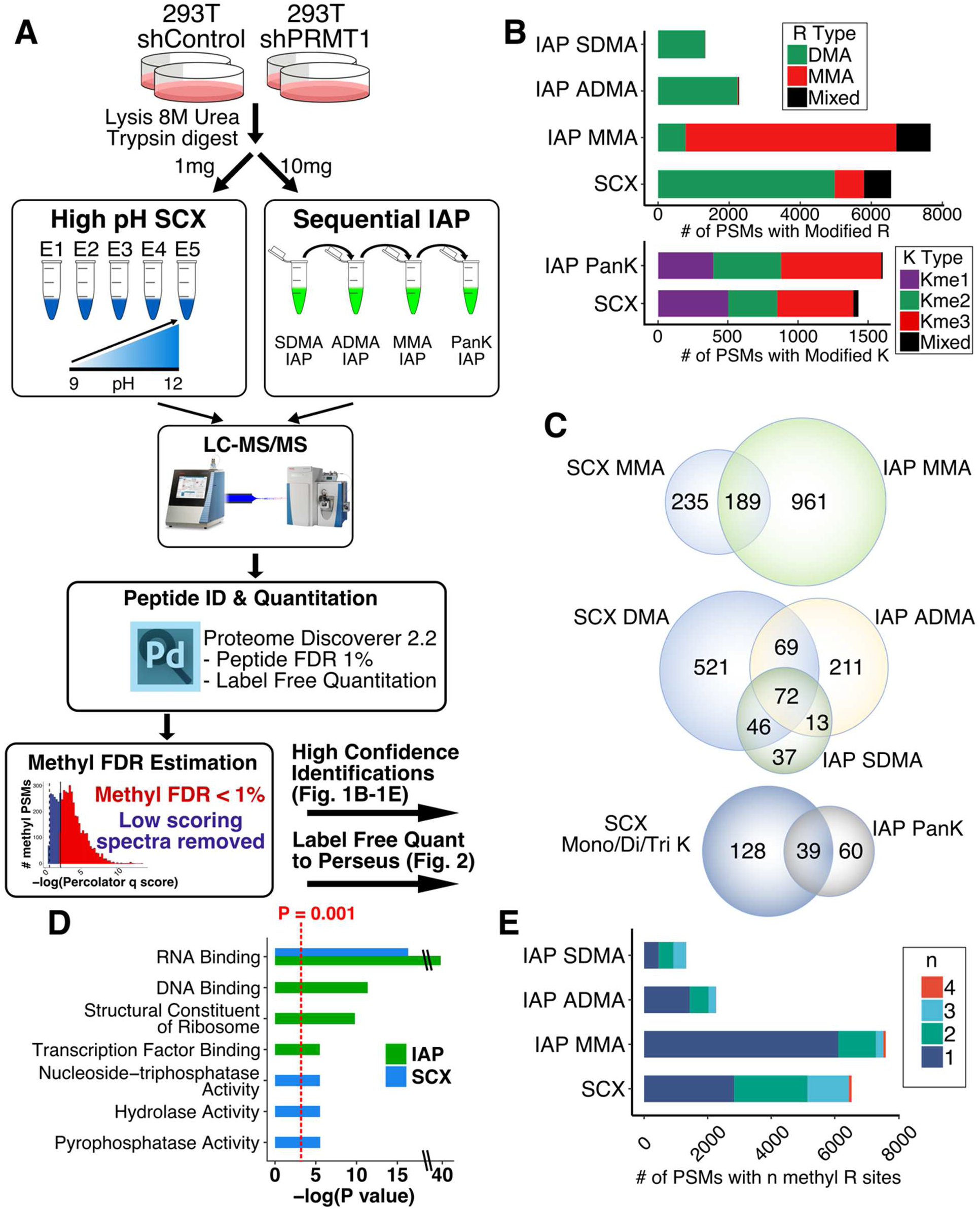
SCX and IAP enrich methyl peptides and target different subsets of protein methylome. A) Schematic of the methyl enrichment workflow for high pH SCX and IAP (21, 28). 293T cells expressing the negative control short hairpin RNA shControl or a short hairpin RNA against PRMT1 (shPRMT1) were lysed in 8 M urea and then digested to peptide with trypsin. Next, 1 mg and 10 mg of tryptic peptides were subjected to high pH SCX or IAP enrichment. For SCX, five fractions eluted at increasing pH were collected. For IAP, the lysates were sequentially incubated with the indicated IAP antibodies. Samples were then analyzed using a Q Exactive Plus mass spectrometer. LC-MS data was searched using Proteome Discoverer, and methyl peptides were subjected to a strict 1% methyl FDR. The number of methyl PSMs for each sample is shown in Table 1. B) Number of methyl PSMs showing the indicated type of arginine methylation for each enrichment technique. Only high confidence spectra passing the 1% methyl FDR were considered. “Mixed” peptides contained a mixture of mono/di methylation on R and mono/di/tri methylation on K on the same peptide. C) SCX and IAP identify different subsets of the protein methylome. Overlap of (top) identified MMA peptides comparing SCX and MMA IAP, (middle) identified DMA peptides comparing SCX and ADMA/SDMA IAPs, and (bottom) identified mono/di/tri-methyl lysine peptides comparing SCX and PanK IAP. D) Gene ontology of methyl peptides enriched by SCX and IAPs. Unique non-overlapping gene symbols from SCX and IAP were compared against a human background using GOrilla. All shown ontologies had an FDR q-value < 0.01 as calculated by GOrilla (59). E) Number of methylation sites per PSM for each methyl peptide enrichment protocol. All IAP methods primarily enriched single methylated peptides, whereas SCX identified a larger fraction of di-, tri-, and tetra-methylated peptides.

**Table 1:** Summmary of Methyl PSMs from SCX and IAP. Summary of spectra identified by each technique. For SCX each of the five fractions are shown along with each IAP. The number of methyl and nonmethyl PSMs were used to calculate the percent enrichment for each fraction and IAP. The number of MMA, Kme1, DMA, Kme2, Kme3, and mixed PSMs are shown. Mixed PSMs contained a mixture of methyl marks on the same peptide (e.g., MMA and DMA). The percolator q-value cutoff used to estimate the methyl FDR is also shown for each technique.

| Strategy | SCX<br>(sum of n = 4 RAW files per SCX fraction) |  |  |  |  |  | IAP<br>(sum of n = 4 RAW files per IAP) |  |  |  |
| --- | --- | --- | --- | --- | --- | --- | --- | --- | --- | --- |
|  | E1 | E2 | E3 | E4 | E5 | All Fxns | SDMA | ADMA | MMA | PanK |
| # Methyl PSMs | 974 | 3326 | 2170 | 1236 | 273 | 7979 | 1332 | 2279 | 7698 | 1605 |
| # Nonmethyl PSMs | 95312 | 61701 | 16460 | 2764 | 1270 | 177507 | 25419 | 19828 | 39194 | 35704 |
| % Enriched | 1.0% | 5.1% | 12% | 31% | 18% | 4.2% | 5.0% | 10% | 16% | 4.3% |
| # PSMs MMA, Kme1 | 146,136 | 242,395 | 207,88 | 71,33 | 5,3 | 671,655 | 4,0 | 20,0 | 5931,15 | 0,394 |
| # PSMs DMA, Kme2 | 313,110 | 2024,200 | 1556,13 | 873,21 | 212,0 | 4978,344 | 1319,3 | 2239,6 | 778,14 | 4,482 |
| # PSMs Kme3 | 241 | 176 | 44 | 74 | 10 | 545 | 0 | 0 | 0 | 717 |
| # PSMs Mixed | 28 | 289 | 262 | 164 | 43 | 786 | 6 | 14 | 960 | 8 |
| # Unique Methyl PSMs | 271 | 798 | 469 | 290 | 94 | 1260 | 194 | 428 | 1456 | 109 |
| Percolator q cutoff<br>Meth FDR% | q < 0.00025<br>1.0% |  |  |  |  |  | q < 0.0017<br>0.90% | q < 0.0027<br>0.87% | q < 0.0027 1.0% | q < 0.0014 1.0% |

### Label free quantitation of methyl peptides shows high reproducibility

Next, we sought to test the reproducibility of SCX and IAP methyl peptide quantitation in 293T cells. Label-free quantitation (LFQ) values of methyl peptides passing the 1% methyl FDR were filtered to remove peptides with missing values, log_2_ transformed, and then normalized by the sample median (Fig. 2A). Across all experiments, we quantified 943 methylation sites on 451 proteins (46% of all identified peptides), of which 262 sites were measured by 2 or more techniques (Tables S8-S15). Similar to identified methyl peptides (Fig. 1C), the overlap between quantified methyl peptides was low between SCX and IAP (Fig. 2B). Scatter plots of LFQ values from biological replicates showed high correlation for each methyl-peptide enrichment technique (Pearson correlation coefficients ranging from 0.85 to 0.95). Together, this data demonstrates that SCX and IAP both generate reproducible LFQ data for methylated peptides.

**Figure 2:**
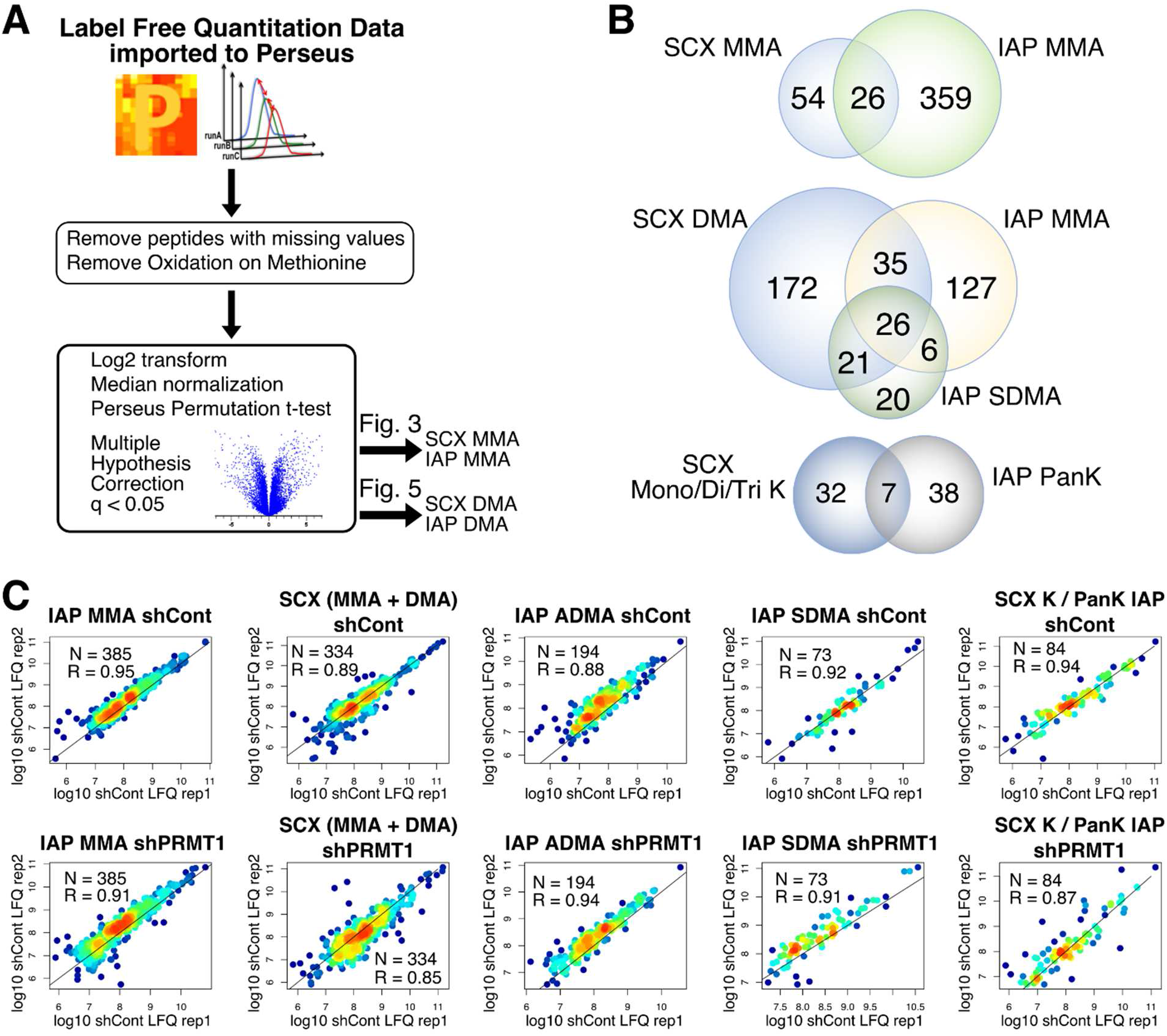
Label free quantitation of methyl peptides is highly reproducible. A) Schematic of the label free quantitation (LFQ) workflow for methyl peptides. LFQ values were imported to Perseus where peptides with missing values or methionine oxidation were removed. Values were log_2_ transformed and median normalized. A permutation-based t-test with an FDR q-value of 0.05 and S_0_ value of 0.5 was performed to identify significantly changing methyl peptides. B) SCX and IAP quantify different subsets of the protein methylome. Overlap of (top) quantified MMA peptides comparing SCX and MMA IAP, (middle) quantified DMA peptides comparing SCX and ADMA/SDMA IAPs, and (bottom) quantified mono/di/tri-methyl lysine peptides comparing SCX and PanK IAP. C) Scatter plots of biological replicates demonstrate high reproducibility. The log10 LFQ values between biological replicates were plotted for each methyl-peptide enrichment. Each dot is colored by the local density of points. The number of peptides and the Pearson correlation coefficient for each comparison is shown.

### Quantitative Analysis of MMA peptides from shPRMT1 293T cells

PRMT1 has been reported to account for over 90% of ADMA methylation events in mammalian cells (48). We therefore investigated how PRMT1 knockdown affected MMA in 293T cells. Knockdown of PRMT1 was confirmed by Western blotting (Fig. 3A). We also observed a general increase in MMA levels in PRMT1 knockdown cells, consistent with other reports (49). SCX and IAP profiling by LC-MS revealed many changing MMA peptides, with good agreement for peptides captured by both techniques (bolded sites, Fig. 3B, Tables S8, S10). Comparing shControl and shPRMT1 cells using a permutation-based t-test in Perseus, we found 61 significantly increased and 58 significantly decreased MMA peptides (q < 0.05) in PRMT1 knockdown cells (Fig. 3C, Table S10). Significantly changing MMA sites were enriched for PRMT1 interactors from the EBI database (Fisher’s Exact p-value = 0.044). Of the total 119 significantly changing methyl peptides, 5 came from SCX enrichment, and 114 from IAP enrichment. Motif analysis of MMA sites with a log_2_ fold change greater than 1.5 in either direction compared against non-significantly changing MMA sites recapitulated the RGG motif common to methyl arginine sites (Fig. 3D). A preference for serine in the −4 and −2 positions and tyrosine in the +3 position was also observed for changing MMA sites. Gene ontology analysis revealed that significantly changing MMA peptides were enriched for mRNA metabolic process and mRNA splicing compared to non-changing MMA peptides (Fig. 3E). Taken together, this data reveals that PRMT1 knockdown dramatically reshaped the MMA proteome.

**Figure 3:**
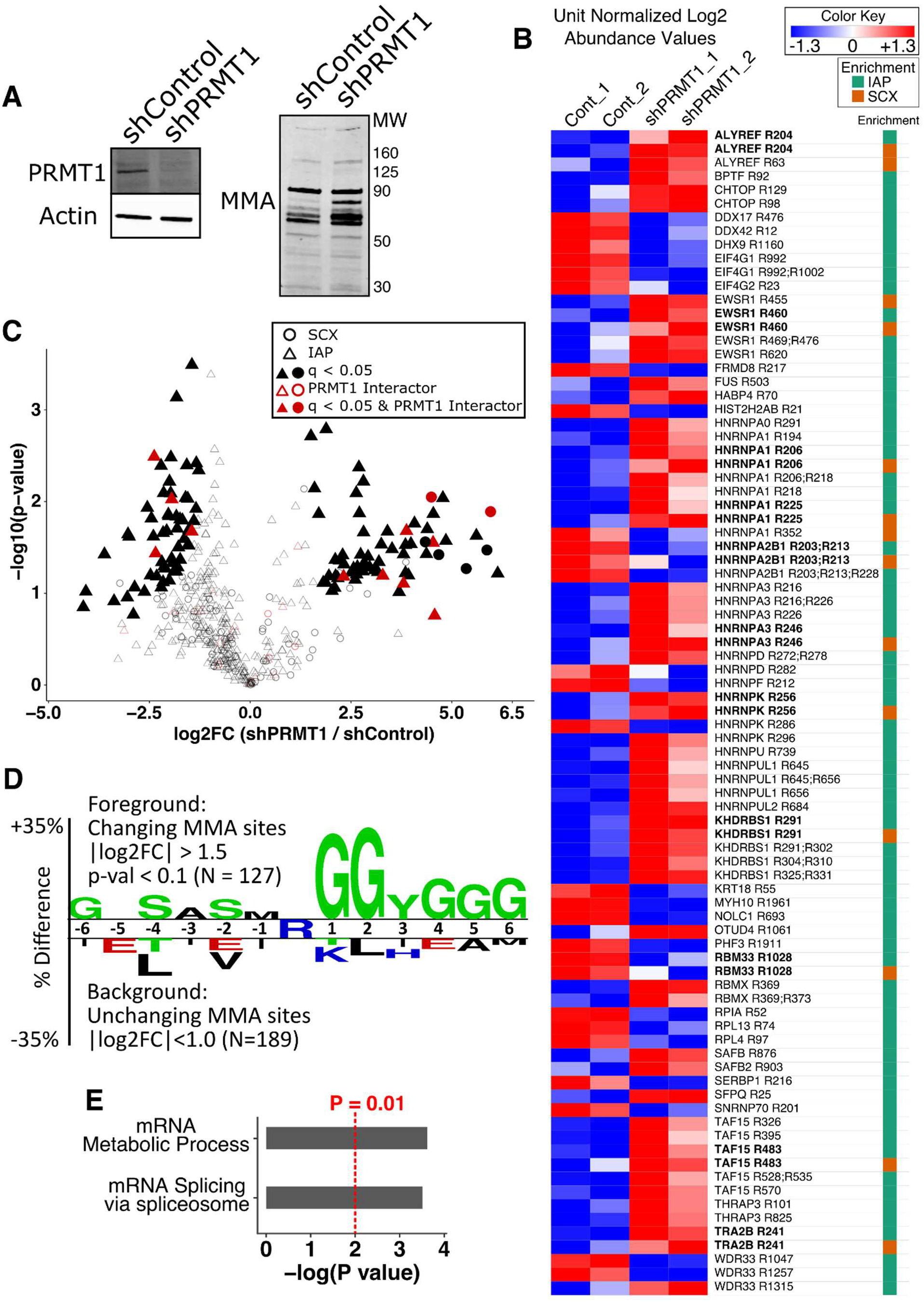
Quantitative Analysis of MMA peptides from shPRMT1 293T cells. A) Western blotting confirmed reduced PRMT1 expression and increased MMA levels upon PRMT1 knockdown. 293T cells expressing shControl or shPRMT1 were lysed and analyzed by Western blotting with antibodies against either PRMT1 or MMA. Actin was used as an equal loading control. B) The MMA methylome is substantially altered by PRMT1 knockdown. Heatmap of peptide level differences for methyl peptides captured by SCX and IAP, sorted by gene name. Median normalized log_2_ LFQ values were unit normalized and colored by fold change as indicated. Methyl peptides measured by both techniques are bolded and showed good quantitative agreement. The column at right indicates which methyl peptide enrichment protocol identified each peptide (SCX or MMA IAP). Bold text indicates peptides identified in both SCX and MMA IAP. C) Volcano plot of MMA peptides enriched by SCX and IAP demonstrating 61 and 58 significantly increased and decreased methyl peptides, respectively, in PRMT1 knockdown cells. The shape indicates which methyl peptide enrichment protocol identified each peptide. Filled shapes indicate q-value < 0.05 by permutation t-test in Perseus. Red points denote known interactors of PRMT1 according to the EBI Int Act database (60), and significantly changing MMA peptides were enriched for PRMT1 interactors (p = 0.044 by Fisher’s Exact test). D) Two sample motif analysis of changing MMA sites recapitulated the known RGG motif of protein arginine methylation. The motif was generated using Two Sample Logo by comparing MMA peptides with absolute value of log_2_ fold change > 1.5 against MMA peptides with absolute value of log_2_ fold change < 1. A p-value of 0.05 was used to generate the motif. E) Gene ontology analysis of MMA peptides with absolute value of log_2_ fold change > 1.5 against MMA peptides with absolute value of log_2_ fold change < 1D) demonstrated that changing MMA sites were enriched for proteins with mRNA metabolic process (GO:0016071) and mRNA splicing (GO:0000398). Enriched ontologies were identified using GOrilla and passed an FDR q-value < 0.25.

### Characteristic neutral losses enable discrimination of SDMA and ADMA

We next sought to investigate the effect of PRMT1 knockdown on protein arginine dimethylation (DMA). Distinguishing the isobaric ADMA and SDMA PTMs should be possible based on characteristic neutral losses from both dimethylarginine forms (26, 32, 50–52). For ADMA, the neutral loss of dimethyamine causes mass loss of 45.058 Da (Fig. 4A), whereas for SDMA, the neutral loss of monomethylamine causes mass loss of 31.042 Da (Fig. 4B). We automated the search for ADMA and SDMA by adding these neutral loss masses to the Andromeda search engine in MaxQuant (53). We then tested the accuracy of this approach using a publicly available data set consisting of synthetic peptides modified with either ADMA or SDMA (50) (Fig. S3) (Pride ID: PXD009449). The synthetic peptides contained identical sequences and a single non-C-terminal ADMA or SDMA. Andromeda successfully identified 54.9% (78/142) of ADMA modifications based on the neutral loss of dimethylamine (Fig. 4C) and 73.1% (106/145) of SDMA modification based on the neutral loss of monomethylamine, with only 1 incorrect identification (i.e., SDMA identified as ADMA). We then applied this analysis to our DMA peptides enriched by SCX and IAP and found between 15-37% of our quantified peptides were annotated as either ADMA or SDMA (Fig. 4D). Manual inspection of identified ADMA and SDMA peptides confirmed the accuracy of this approach (Fig. 4E,F). Although SCX should equally enrich both ADMA and SDMA, a large majority of neutral loss annotated DMA peptides (86%, 228 of 265 total) were annotated as ADMA. In addition, we found that the IAP antibodies exhibited a strong preference for their intended targets, with 87.5% and 78.4% of quantified peptides identified by ADMA and SDMA IAP identified as ADMA and SDMA, respectively. Thus, the neutral loss of dimethylamine and monomethylamine can unambiguously discriminate ADMA and SDMA, respectively.

**Figure 4:**
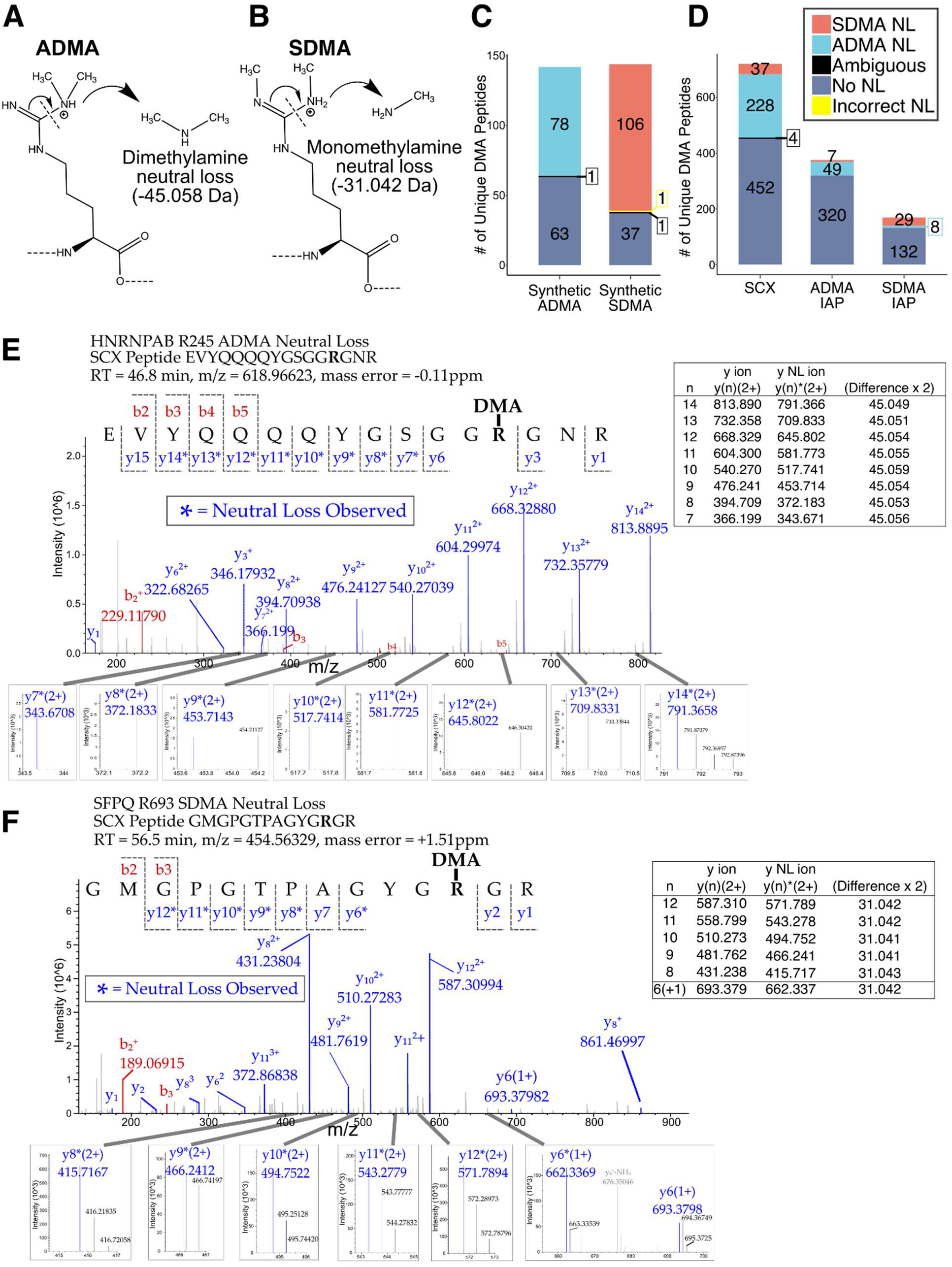
Characteristic neutral loss of methylamine and dimethylamine allows discrimination of SDMA and ADMA spectra respectively. A) Mechanism of neutral loss for ADMA resulting in neutral loss of dimethylamine (45.058 Da). B) Mechanism of neutral loss for SDMA resulting in neutral loss of monomethylamine (31.042 Da). C) Analysis of synthetic ADMA and SDMA peptides from Zolg et al. (50) were searched for neutral losses using the Andromeda search engine. The synthetic peptides contained identical sequences and a single non-C-terminal ADMA or SDMA. Ambiguous identifications were defined as peptides who had conflicting neutral loss assignments and whose mean Andromeda scores differed by less than 30. Andromeda successfully identified 54.9% (78/142) of ADMA modifications and 73.1% (106/145) of SDMA modifications. Only one SDMA peptide was incorrectly identified as ADMA. D) MaxQuant neutral loss search applied to quantified DMA peptides from 293T cells expressing either shControl or shPRMT1 after enrichment by either SCX or ADMA/SDMA IAPs. Peptides showing neutral loss from the Andromeda search in MaxQuant were matched to their corresponding peptides in the LFQ data from Proteome Discoverer 2.2. Matching was performed by matching peptide sequence, retention time, methyl sites, and sample origin between neutral loss data and LFQ data. E) Annotated spectra of an ADMA peptide from SCX showing neutral losses of 45.058 for nine y ions. The inset peaks of each neutral loss fragment are shown with some fragments showing the isotopic envelope typical of peptides. An inset table shows the masses for each y ion and y ion neutral loss fragment, as well as the differences in mass for each pair. F) Annotated spectra of an SDMA peptide from SCX showing neutral losses of 31.042 Da for six y ions. The inset peaks of each neutral loss fragment are shown with some fragments showing the isotopic envelope typical of peptides. An inset table shows the masses for each y ion and y ion neutral loss fragment, as well as the differences in mass for each pair.

### Quantitative Analysis of DMA peptides from shPRMT1 293T cells

We next investigated how PRMT1 knockdown affected ADMA and SDMA in 293T cells. Immunoblotting for ADMA and SDMA on shPRMT1 293T lysates revealed a slight decrease in ADMA methylation and an increase in SDMA methylation in PRMT1 knockdown cells (Fig. 5A). SCX and IAP profiling by LC-MS revealed many significantly changing DMA peptides, with good quantitative agreement for DMA peptides identified by both SCX and IAP (bolded sites, Fig. 5B Tables S9, S11-12). For each methyl enrichment technique (SCX, ADMA IAP, SDMA IAP), we compared shControl and shPRMT1 cells using a permutation-based t-test in Perseus. We found 2 significantly increased SCX peptides (Fig. 5C), 4 significantly increased ADMA IAP peptides (Fig. 5D), and 3 significantly increased SDMA IAP peptides (Fig. 5E) (q < 0.05). Two methyl peptides (DHX9 R1249/R1253/R1265 and HNRNPA3 R246) were significantly upregulated in both ADMA and SDMA IAP data sets. SCX also enriched peptides with both MMA and DMA modifications, creating “mixed” methyl peptides, including four significantly changing peptides (Fig. S4, Table S15). Motif analysis of the downregulated ADMA peptides revealed a preference for arginine in the −2 position and leucine/aspartic acid/asparagine in the −1 position (Fig. 5F). Gene ontology analysis revealed that significantly changing ADMA peptides were enriched for nitrogen compound transport, organic substance transport, RNA localization, and protein localization compared to the background (Fig. 5G).

**Figure 5:**
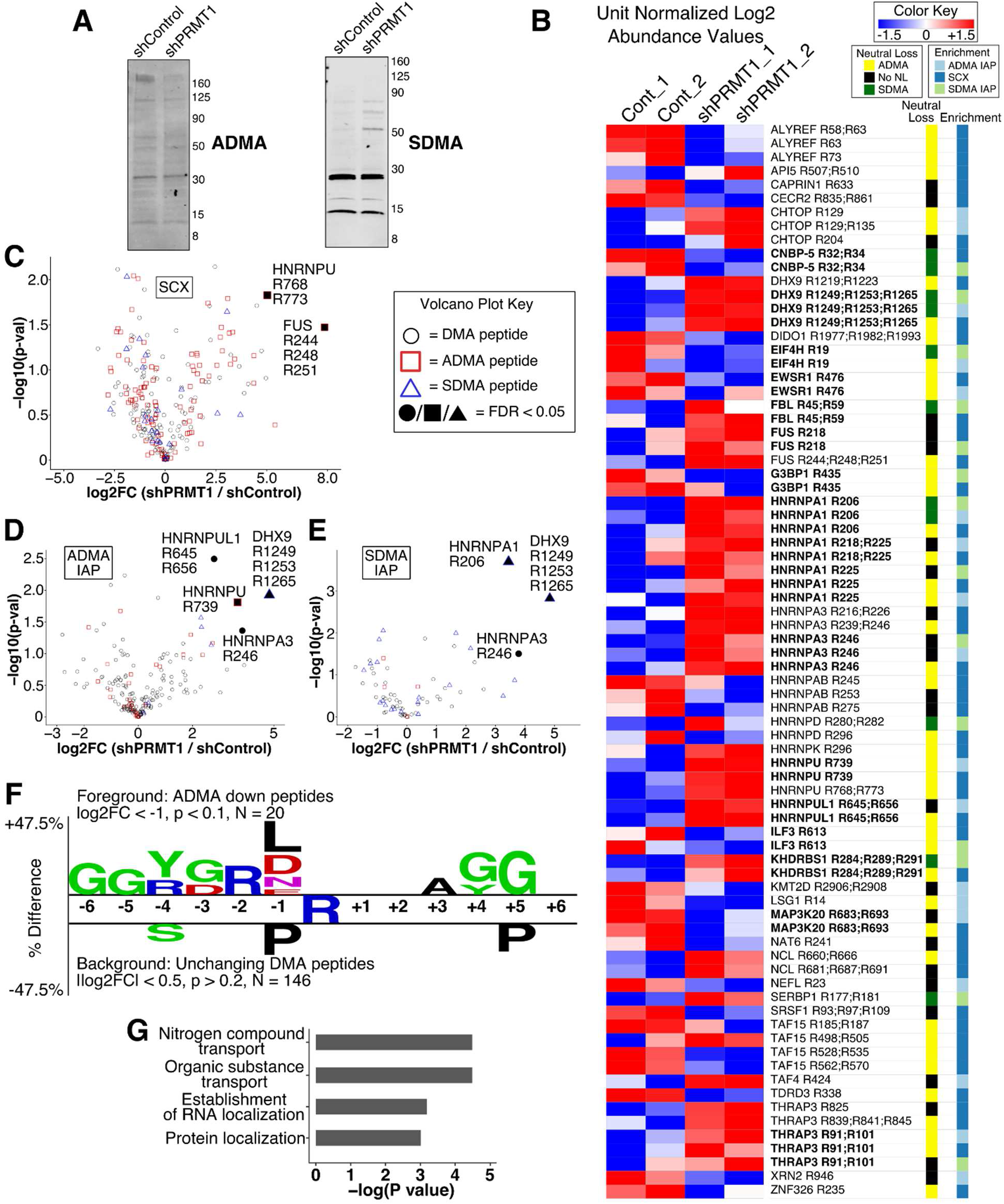
Quantitative Analysis of DMA peptides from shPRMT1 293T cells. A) Immunoblot of ADMA and SDMA on 293T cells expressing shControl or shPRMT1. 293T cells expressing shControl or shPRMT1 were lysed and analyzed by Western blotting with antibodies against either ADMA or SDMA. ADMA levels were decreased, and SDMA levels were increased in PRMT1 knockdown cells. Actin was used as an equal loading control and is shown in Fig. 3A. B) The DMA methylome is substantially altered by PRMT1 knockdown. Heatmap of dimethyl peptide levels enriched by SCX, ADMA IAP, or SDMA IAP, sorted by gene name. Median normalized log_2_ LFQ values were unit normalized and colored by fold change as indicated. The columns at right indicate observed neutral losses (ADMA or SDMA) and which methyl peptide enrichment protocol identified each peptide. Bold text indicates peptides that were identified by multiple enrichment protocols. C-E) Volcano plots of DMA peptides enriched by SCX (C), ADMA IAP (D), and SDMA IAP (E) demonstrating significantly increased DMA peptides in PRMT1 knockdown cells. The type of neutral loss, if observed, is indicated by shape, and the filled in points indicate q-value < 0.05 by permutation t-test in Perseus. F) Two-sample motif analysis of downregulated dimethyl peptides showing ADMA neutral loss compared to unchanging background dimethyl peptides. All downregulated ADMA peptides from all experiments were combined for the foreground with log_2_ fold change < −1 and a p-value < 0.1. All unchanging DMA peptides from all experiments were used as the background with a log_2_ fold change between 0.5 and −0.5 with a p-value > 0.20. G) Gene ontology analysis revealed that proteins with significantly changing ADMA sites were enriched for nitrogen compound transport, organic substance transport, establishment of RNA localization, and protein localization. ADMA peptides with absolute value log_2_ fold change > 1 and Student’s t-test p-value < 0.1 were compared against ADMA peptides with absolute value log_2_ fold change < 0.5 and Student’s t-test p-value > 0.2. Enriched ontologies were identified in GOrilla and passed an FDR q-value < 0.25.

### Lysine methylation is not affected by PRMT1 knockdown

In addition to arginine methylation, both SCX and IAP can enrich peptides containing methyl lysine. To test whether PRMT1 depletion affected protein lysine methylation, 293T cells expressing shPRMT1 or the shControl were subjected to both SCX and Pan-methyl-K IAP followed by mass spectrometry. Label-free quantitation of mono-, di-, and tri-methyl lysine by IAP showed only 1 site significantly changed for SCX (HMGN2 K40, Fig. S5A, Table S13), and no significant changes in protein lysine methylation for PanK IAP (Fig. S5B, Table S14). Together, these data demonstrate that PRMT1 depletion affects methylation of arginine but not lysine residues.

### Integrated analysis of methyl-arginine forms reveals novel PRMT1 substrates and substrate scavenging

Because PRMT1 catalyzes both MMA and ADMA, PRMT1 substrates may exhibit both downregulated MMA and ADMA levels in PRMT1 knockdown cells (Fig. 6A). However, because other PRMT1s can also catalyze MMA, it is possible that PRMT1 targets will exhibit downregulated ADMA but upregulated MMA levels. We therefore reasoned that integrating results from MMA and DMA would enable a more comprehensive view of the PRMT1 methylome. We identified 17 methylation sites on 11 proteins that exhibited downregulated DMA levels and confirmed ADMA neutral loss (Fig. 6B). Several of these methylation sites exhibited increased MMA levels concomitant with decreased ADMA levels (e.g., EWSR1 R460). In addition, we found 12 methylation sites on 9 proteins with upregulated DMA levels and confirmed ADMA neutral loss (Fig. 6C), consistent with scavenging by other Type 1 PRMTs in the absence of PRMT1 activity.

**Figure 6:**
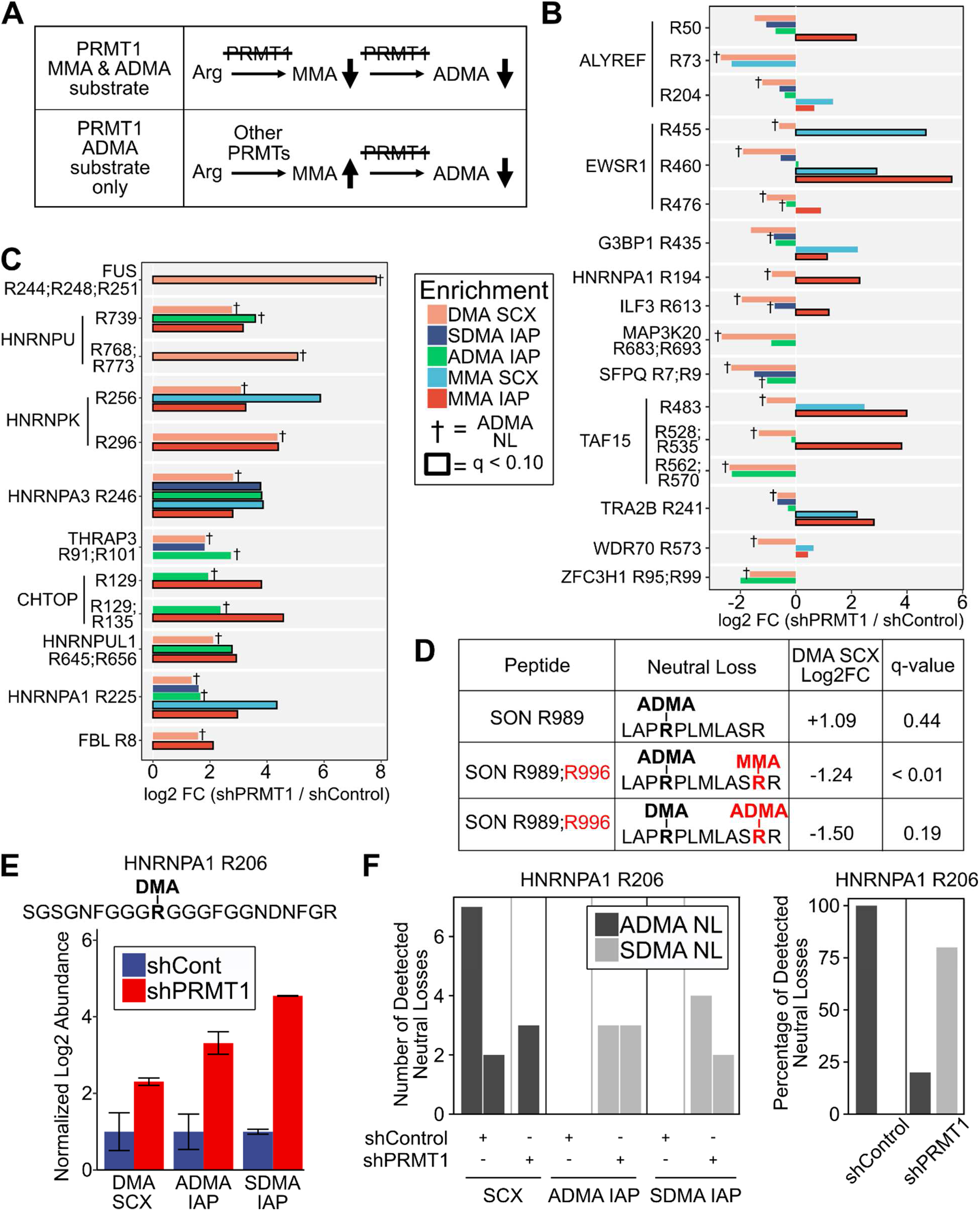
Integrated analysis of methyl-arginine forms reveals novel PRMT1 substrates and ADMA substrate scavenging. A) Schematic depicting the expected trends in MMA and ADMA levels for methylation sites targeted by PRMT1 for both MMA and ADMA methylation (top) and ADMA but not MMA methylation (bottom). B) Integrated analysis of MMA and ADMA levels revealed novel PRMT1 substrates. Log_2_ fold change of methylation levels for shPRMT1 cells compared to shControl cells are shown for different methyl peptide enrichment protocols: DMA peptides identified by SCX, SDMA IAP, ADMA IAP, MMA peptides identified by SCX, and MMA IAP. Peptides were selected based on decreased ADMA levels in one or more experiments, and the presence of MMA data for the same methylation site. † denotes methyl peptides with confirmed ADMA neutral loss. Bold outline indicates methyl peptides with FDR q-value < 0.1 by permutation t-test in Perseus. C) Integrated analysis of MMA and ADMA levels revealed substrate scavenging in the absence of PRMT1 activity. Log_2_ fold change of methylation levels for shPRMT1 cells compared to shControl cells are shown for different methyl peptide enrichment protocols: DMA peptides identified by SCX, SDMA IAP, ADMA IAP, MMA peptides identified by SCX, and MMA IAP. Peptides were selected based on increased ADMA levels in one or more experiments, and the presence of MMA data for the same methylation site. † denotes methyl peptides with confirmed ADMA neutral loss. Bold outline indicates methyl peptides with FDR q-value < 0.1 by permutation t-test in Perseus. D) Identification of SON R996 as a PRMT1 substrate. A peptide with neutral loss confirmed ADMA R989 was upregulated upon PRMT1 knockdown. A peptide with neutral loss confirmed ADMA R989 and MMA R996 was downregulated significantly in the mixed SCX dataset. A peptide with DMA R989, neutral loss confirmed ADMA R996, and a missed cleavage at R996 was downregulated upon PRMT1 knockdown. This suggests that PRMT1 knockdown reduced ADMA R996 and MMA R996, allowing tryptic cleavage, thereby resulting in increased levels of the fully cleaved tryptic peptide with ADMA R989. E-F) HNRNPA1 R206 exists in both ADMA and SDMA modified form and may switch from ADMA to SDMA upon PRMT1 knockdown. (E) The methyl peptide SGSGNFGGG<u>*R*</u>GGGFGGNDNFGR (DMA site underlined and italicized) was upregulated in SCX, ADMA IAP, and SDMA IAP. The log_2_ fold change of shPRMT1 cells compared to shControl cells is shown, normalized to shControl. (F) Analysis of neutral loss ions demonstrated that ADMA neutral losses were primarily identified in shControl cells, whereas SDMA neutral losses were primarily identified in shPRMT1 cells. (left) Each bar represents a PSM with the y-axis representing the number of neutral losses observed for that PSM. There were no identified neutral losses in either ADMA IAP or SDMA IAP in the shControl cells. (right) The percentage of ADMA and SDMA neutral losses observed for shControl and shPRMT1 cells.

Next, because arginine dimethylation is more likely to result in missed tryptic cleavages (28), we reasoned that we might identify PRMT1 substrates by examination of ADMA peptides with and without missed tryptic cleavages (Fig. S6). Using this approach, we identified SON R996 as a high confidence PRMT1 target. In our SCX data, SON R996 was present in three forms: unmethylated with tryptic cleavage at R996 (i.e., LAPRPLMLAS**<u>R</u>**, R996 in bold underline), monomethylated with missed cleavage at R996 (i.e., LAPRPLMLAS**<u>R</u>**R), and ADMA-modified with missed cleavage at R996 (i.e., LAPRPLMLAS**<u>R</u>**R). The levels of the unmethylated, MMA, and ADMA peptides were increased (log_2_ fold change +1.09), decreased (log_2_ fold change −1.24), and decreased (log_2_ fold change −1.50), respectively, in PRMT1 knockdown cells (Fig. 6D). In total, we found 4 examples where identification of peptides with and without missed cleavages enabled deeper understanding of methylation dynamics, although SON R996 was the only putative PRMT1 target (Fig. S6). Finally, we identified one DMA site, HNRNPA1 R206, that was increased in abundance in SCX, ADMA IAP, and SDMA IAP data (Fig. 6E) and that exhibited both ADMA and SDMA neutral losses. For this methyl peptide, ADMA neutral losses were more frequent in shControl cells, and SDMA neutral losses were more prevalent in shPRMT1 cells (Fig. 6F).

## DISCUSSION

Despite its relevance for signal transduction, metabolism, transcription, and other cellular phenotypes, protein arginine and lysine methylation remain understudied. This is partly because of the inherent difficulty of enriching a small, neutral PTM and partly because methyl peptide enrichment strategies have been less comprehensively studied than other PTMs. In this study, we compared two methyl peptide enrichment techniques: high pH SCX and IAP (Figs. 1, 2). We found that the two techniques were mostly orthogonal for both methyl peptide identification and LFQ quantitation, demonstrating that comprehensive measurement of the protein methylome requires multiple methyl peptide enrichment strategies. Although SCX and IAP enrich different methyl peptides, both SCX and IAP peptides were enriched for the GO annotation RNA binding, consistent with the known function of protein methylation (11, 24, 43–47) (Fig. 1D). One explanation for the low overlap between SCX and IAP is the tendency of DMA to result in missed tryptic cleavages (28). Because SCX enriches highly positively charged peptides, SCX preferentially identifies multi-methylated peptides with missed cleavages and therefore more positive charge (Fig. 1E). In contrast, because MMA is readily cleaved by trypsin, MMA IAP enriches significantly more MMA peptides than SCX because these peptides are less likely to have missed cleavages.

Using these orthogonal methyl peptide enrichment techniques, we investigated how PRMT1 knockdown remodeled the protein methylome and found significant changes to 127 methylarginine sites (q < 0.05) on 78 proteins (Fig. 3, 5). We observed that PRMT1 knockdown significantly affected only one lysine methylation site (Fig. S5), although our data support previous observations that lysine methylation is much less abundant *in vivo* than arginine methylation (21, 28). Of the significantly changing arginine methylation sites, the large majority were MMA (119 of 127, or 93.7%), with most significantly changing MMA sites identified by IAP rather than SCX (114 of 119, or 95.8%). Because PRMT1 catalyzes MMA modifications, the 58 significantly downregulated MMA sites we observed in PRMT1 knockdown cells may represent PRMT1 MMA targets (Fig. 3C). Conversely, because accumulation of MMA can result from inhibition of PRMT1-mediated ADMA modification, the 61 significantly increased MMA sites we observed in PRMT1 knockdown cells may represent PRMT1 ADMA sites. Consistent with this hypothesis, significantly changing MMA sites were enriched for known PRMT1 interactors (Fisher’s Exact p-value = 0.044). In addition, identification of the RGG motif from significantly changing MMA sites (Fig. 3D) further confirmed that PRMT1 targets GAR (glycine arginine rich) motifs (54). Taken together, these results demonstrate the PRMT1-mediated regulation of MMA in 293T cells.

Recent reports have indicated that the neutral losses of dimethylamine and methylamine can discriminate between ADMA and SDMA, respectively (26, 32, 50–52). Here, we have extended those findings by reanalyzing data from synthetic peptides with either ADMA or SDMA modifications (50). Although 25-45% of synthetic peptides did not generate identifiable neutral losses (Fig. 4C), the accuracy of ADMA and SDMA identification was very high for spectra with identified neutral losses: 78/78 and 106/107 peptides for ADMA and SDMA, respectively. We subsequently applied this approach to both SCX and IAP methyl peptide enrichment strategies. In addition to confirming the general specificity of the ADMA and SDMA IAP antibodies, we found that SCX, a technique which should not be biased towards either ADMA or SDMA peptides, identified 228 ADMA but only 37 SDMA peptides (Fig. 4D). This result indicates ADMA may be more prevalent in mammalian cells than SDMA (6:1 ratio), consistent with observations in mouse embryonic fibroblasts that SDMA is present in proteins at 10-fold lower concentration than ADMA (49).

Analysis of arginine dimethylation also revealed considerable changes upon knockdown of PRMT1. In PRMT1 knockdown cells, we observed decreased ADMA levels by immunoblot, consistent with loss of PRMT1’s type I methyltransferase activity (Fig. 5A). Additionally, we observed increased SDMA levels by immunoblot in PRMT1 knockdown cells, consistent with substrate scavenging by type II PRMTs in the absence of PRMT1 activity. In our LC-MS data, we identified several candidates for type II PRMT scavenging including HNRNPA1 206, FBL R45;R49, and SERBP1 R177;R181 (Fig. 5B). Interestingly, HNRNPA1 showed evidence of methyl switching from ADMA to SDMA. First, levels of HNRNPA1 R206 were increased in PRMT1 knockdown cells in SCX, ADMA IAP, and SDMA IAP data sets (Fig. 6E). Second, analysis of the neutral losses in each data set revealed PRMT1 knockdown increased the percentage of SDMA neutral losses (0% in shControl and 80% in shPRMT1 cells) and decreased the percentage of ADMA neutral losses (100% in shControl and 20% in shPRMT1 cells) (Fig. 6F). Taken together, this data indicates that HNRNPA1 R206 can exist as either ADMA and SDMA and suggests that PRMT1 knockdown results in a switch from ADMA to SDMA. This finding is particularly interesting because switching between methyl forms can affect protein function (12) and PRMT5-mediated methylation of HNRNPA1 regulates translation mRNAs containing internal ribosome entry sites (44). Thus, PRMT1 and the interplay between ADMA and SDMA modifications may regulate HNRNPA1 activity.

Furthermore, we identified high confidence PRMT1 ADMA targets by integrating different forms of arginine methylation (eg. MMA, SDMA, ADMA) (Fig. 6A). For example, PRMT1 knockdown resulted in decreased ADMA levels at EWSR1 R460 (log_2_ fold change −1.91, q-value 0.14 in SCX DMA with ADMA neutral loss), suggesting that R460 may be a PRMT1 ADMA target. However, by considering that PRMT1 knockdown also significantly increased the levels of EWSR1 R460 MMA (log_2_ fold change 2.91, q-value 0.09 in SCX MMA and log_2_ fold change 5.61, q-value < 0.004 in MMA IAP), our confidence that EWSR1 R460 is a PRMT1 ADMA site is increased. Using this approach, we identified 17 high confidence PRMT1 substrates (Fig. 6B). In addition, we identified one additional high confidence PRMT1 target by comparing peptides with and without missed cleavages. We identified three peptides with and without missed tryptic cleavages from the protein SON that contained R996 either without methylation or with MMA or ADMA modifications (Fig. 6D). Because PRMT1 resulted in increased levels of the unmethylated peptide and decreased levels of both the MMA and ADMA peptides, this data suggests that SON R996 is both a target of PRMT1 MMA and ADMA modification. A similar integrative analysis yielded ADMA sites that we predict are scavenged by other type I PRMTs in the absence of PRMT1 (Fig. 6B). These peptides all contained ADMA neutral loss and showed increased ADMA and MMA levels in PRMT1 knockdown cells. However, for these peptides, we cannot exclude the possibility that increased methylation is driven by increases in total protein abundance. Taken together, our data demonstrate the value of integrative analysis to elucidate the complex dynamics of arginine methylation.

Taken together, our data highlight that PRMT1 is likely to regulate both RNA:protein interactions and the protein subcellular localization. Although methylated proteins are generally enriched for RNA binding proteins (Fig. 1D), both significantly changing MMA sites and significantly decreased ADMA sites in PRMT1 knockdown cells were further enriched for RNA-related processes (Figs. 3E and 5G). In addition, of the 12 proteins on which we identified 18 high confidence PRMT1 targets, 10 are known RNA binding proteins (i.e., all proteins in Figs. 6B and D except MAP3K20 and WDR70). In addition, arginine methylation can affect protein subcellular localization, including the nucleo-cytoplasmic shuttling of RNA binding proteins (55, 56). In our data, PRMT1 knockdown decreased MMA of DHX9 R1160, a residue that regulates the nuclear localization of DHX9 (57). Interestingly, SDMA levels of DHX9 on the neighboring residues R1249/R1253/R1265 were increased in PRMT1 knockdown cells, suggesting that DHX9 becomes more accessible to Type II PRMTs when localized to the cytoplasm.

In summary, our results confirm that PRMT1 regulates a substantial amount of arginine methylation in mammalian cells. The fact that over 90% of significantly changing methyl arginine sites are not known interactors of PRMT1 demonstrates the need for continued comprehensive analysis of PRMTs and their substrates. This is especially relevant considering the growing body of evidence that dysregulation of arginine methylation may contribute to diseases affecting cancer (14). Our findings validate the utility of using high pH SCX and IAP for enrichment of methyl peptides and to enhance coverage and quantitation of the methylome. The dynamic interplay between different methylation marks highlights the need for further development of methods to quantify site occupancy across all methylation forms, as has been done for simpler PTMs including phosphorylation (58). Towards this end, improved methods to distinguish ADMA and SDMA through fragmentation patterns will be valuable. Finally, given our demonstration that high pH SCX and IAP are largely orthogonal, the continued incorporation of fractionation techniques (24) and alternative methyl-peptide enrichment strategies (29–31) will enable deeper analysis of the protein methylome.

## Supporting information

Supplemental Tables Part 2

Supplemental Tables Part 1

## ACKNOWLEDGMENTS

This work was supported by the Viterbi School of Engineering (NAG), grant R01-DE026468 from the National Institute of Dental and Craniofacial Research (JX), and a Research Career Development Award from STOP CANCER Foundation (JX).

## DATA AVAILABILITY

The mass spectrometry proteomics data have been deposited to the ProteomeXchange Consortium (http://proteomecentral.proteomexchange.org) via the PRIDE partner repository with the dataset identifier PXD012357 (Username, password: 5WbkZaM5).

