## Supplemental Figures for "Deep protein methylation profiling by combined chemical and immunoaffinity approaches reveals novel PRMT1 targets"

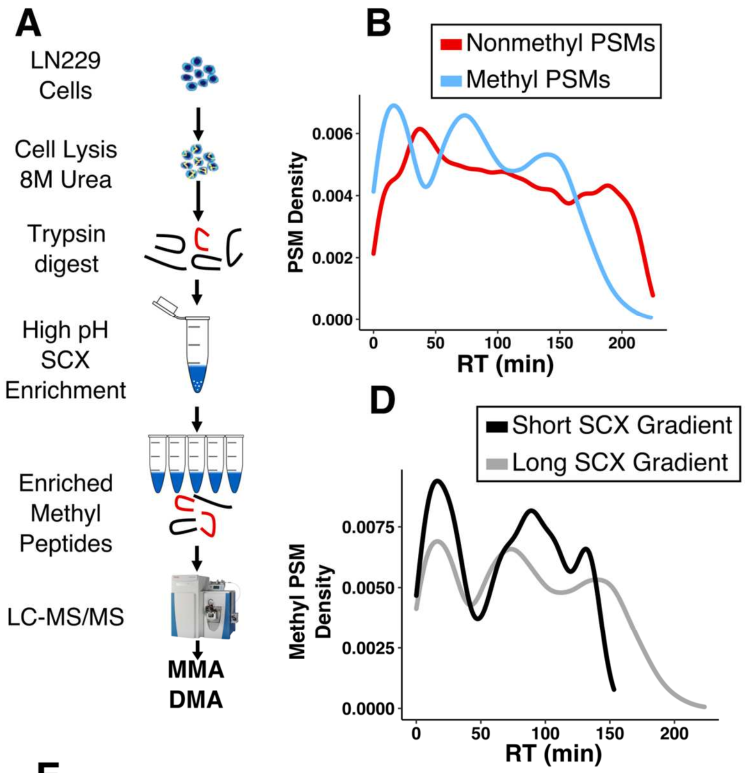


**C**

Long SCX gradient

| Time (min) | Buffer A (%) | Buffer B (%) | |
|---|---|---|---|
| 0 | 98 | 2 | Peptide Loading |
| 20 | 98 | 2 | |
| 25 | 95 | 5 | Peptide Separation |
| 175 | 75 | 25 | |
| 205 | 55 | 45 | |
| 215 | 10 | 90 | Wash |
| 225 | 10 | 90 | |

Short SCX gradient

| Time (min) | Buffer A (%) | Buffer B (%) | |
|---|---|---|---|
| 0 | 100 | 0 | Peptide Separation |
| 50 | 93 | 7 | |
| 142 | 72 | 29 | |
| 152 | 10 | 90 | Wash |
| 155 | 10 | 90 | |

**E**

| Strategy | SCX Long Gradient 225 min (sum of n = 1 RAW file per SCX fraction) | | | | | |
|---|---|---|---|---|---|---|
| Sample | E1 | E2 | E3 | E4 | E5 | All Fxns |
| # Methyl PSMs | 325 | 1214 | 1306 | 502 | 166 | 3513 |
| # Nonmethyl PSMs | 35066 | 18611 | 6263 | 2608 | 1270 | 63818 |
| **% Enriched** | **0.9%** | **6.1%** | **17%** | **16%** | **12%** | **5.2%** |
| # PSMs MMA, Kme1 | 21,40 | 125,50 | 78,82 | 33,15 | 0,1 | 257,188 |
| # PSMs DMA, Kme2 | 115,57 | 828,42 | 906,35 | 376,2 | 156,0 | 2381,136 |
| # PSMs Kme3 | 90 | 33 | 33 | 11 | 1 | 168 |
| # PSMs Mixed | 2 | 126 | 167 | 65 | 8 | 368 |
| # Unique Methyl PSMs | 111 | 353 | 275 | 123 | 48 | 626 |
| Percolator q cutoff Meth FDR% | q < 0.00045 0.96% | | | | | |

| Strategy | SCX Short Gradient 155 min (sum of n = 1 RAW file per SCX fraction) | | | | | |
|---|---|---|---|---|---|---|
| Sample | E1 | E2 | E3 | E4 | E5 | All Fxns |
| # Methyl PSMs | 248 | 765 | 1194 | 450 | 175 | 2832 |
| # Nonmethyl PSMs | 27015 | 17192 | 7196 | 2225 | 1672 | 55300 |
| **% Enriched** | **0.9%** | **4.3%** | **14%** | **17%** | **9.5%** | **4.9%** |
| # PSMs MMA, Kme1 | 20,33 | 93,25 | 127,85 | 25,31 | 5,3 | 270,177 |
| # PSMs DMA, Kme2 | 85,56 | 491,42 | 717,51 | 309,2 | 139,0 | 1741,151 |
| # PSMs Kme3 | 51 | 24 | 57 | 6 | 2 | 140 |
| # PSMs Mixed | 2 | 82 | 156 | 77 | 26 | 343 |
| # Unique Methyl PSMs | 90 | 290 | 272 | 134 | 50 | 550 |
| Percolator q cutoff Meth FDR% | q < 0.00033 0.98% | | | | | |

**Fig. S1. Gradient improvement for high pH SCX enrichment.**

A) Workflow of high pH SCX enrichment (1). LN229 cells were lysed in 8M urea and digested with trypsin. From 1 mg of protein, five fractions are produced by the SCX procedure and were subsequently run on a Q Exactive Plus MS. The data was searched on Proteome Discoverer 2.2 and methyl peptides were subject to a strict 1% methyl FDR. The number of PSMs for two gradients tested is shown in the table in E).

B) Density plots of PSM retention times for methyl and nonmethyl PSMs run on the "Long" SCX gradient used from Wang *et al*.. PSMs from all 5 fractions were combined and their retention times plotted. The average density of nonmethyl PSMs and methyl PSMs was 275 PSM / min and 15 PSMs / min, respectively.

C) Original "Long" gradient and a new proposed "Short" gradient for SCX. Wang *et al*. did not collect spectra for the first 20 min during sample loading, but because we found that methyl PSMs were eluting during this sample loading phase, we did collect spectra during this phase of the LC gradient.

D) Density plot of methyl PSM retention time for methyl PSMs captured by the "Long" and "Short" SCX gradients. PSMs from all 5 fractions were combined and their retention times plotted for each gradient. The average density of "Long" and "Short" gradient methyl PSMs was 15 PSM / min and 19 PSMs / min, respectively.

E) Summary of spectra identified by each gradient. Each of the five SCX fractions are shown individually. The numbers of methyl and nonmethyl PSMs were used to calculate the percent enrichment for each fraction. The number of MMA, Kme1, DMA, Kme2, PSMs, and mixed methyl PSMs are shown. Mixed PSMs contained a mixture of methyl marks on the same peptide (e.g., MMA and DMA). The percolator q-value cutoff used to estimate the methyl FDR is also shown for each technique.

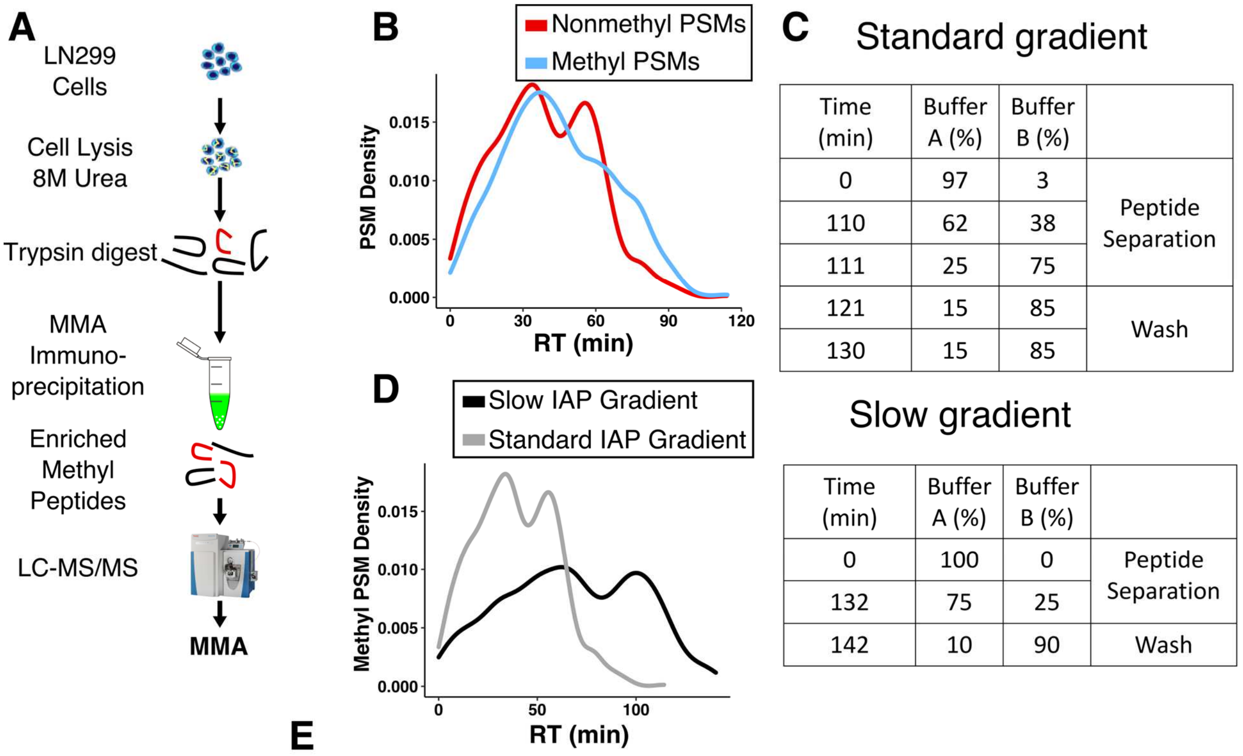


**C — Standard gradient**

| Time (min) | Buffer A (%) | Buffer B (%) | |
|---|---|---|---|
| 0 | 97 | 3 | Peptide Separation |
| 110 | 62 | 38 | |
| 111 | 25 | 75 | |
| 121 | 15 | 85 | Wash |
| 130 | 15 | 85 | |

**Slow gradient**

| Time (min) | Buffer A (%) | Buffer B (%) | |
|---|---|---|---|
| 0 | 100 | 0 | Peptide Separation |
| 132 | 75 | 25 | |
| 142 | 10 | 90 | Wash |

**E**

| Strategy | Standard IAP gradient 130 min (sum of n = 1 RAW file) | Slow IAP gradient 142 min (sum of n = 1 RAW file) |
|---|---|---|
| Sample | **MMA IAP** | **MMA IAP** |
| # Methyl PSMs | 1406 | 1684 |
| # Nonmethyl PSMs | 2709 | 3240 |
| **% Enriched** | **28%** | **34%** |
| # PSMs MMA, Kme1 | 1056,2 | 1205,1 |
| # PSMs DMA, Kme2 | 156,1 | 205,1 |
| # PSMs Kme3 | 0 | 0 |
| # PSMs Mixed | 189 | 272 |
| # Unique Methyl PSMs | 590 | 720 |
| Percolator q cutoff Meth FDR% | q < 0.006 0.99% | q < 0.004 0.83% |

**Fig. S2. Gradient improvement for MMA IAP enrichment.**

A) Workflow of MMA IAP enrichment. LN229 cells were lysed in 8M urea and digested with trypsin. 10 mg of protein were incubated with commercial anti-MMA antibodies conjugated to agarose beads, washed, and eluted, and subsequently run on a Q Exactive Plus MS. The data was searched on Proteome Discoverer 2.2 and methyl peptides were subject to a strict 1% methyl FDR. The number of PSMs for two gradients tested is shown in the table in E).

B) Density plots of PSM retention times for methyl and nonmethyl PSMs run on an in-house "Standard" gradient, similar to other IAP methods (2). The average density of nonmethyl and methyl PSMs was 22 PSM / min and 12 PSMs / min, respectively.

C) In-house "Standard" proteomics gradient and a new proposed "Slow" gradient for MMA IAP. The "Slow" gradient extends the length of the gradient slightly but uses a slower ramp of acetonitrile.

D) Density plot of methyl PSM retention time for methyl PSMs captured by the "Standard" and "Slow" IAP LC gradients. The average density of methyl PSMs for the "Standard" and "Short" LC gradients was 12 PSM / min and 13 PSMs / min, respectively.

E) Summary of spectra identified by each gradient. The numbers of methyl and nonmethyl PSMs were used to calculate the percent enrichment for each fraction and IAP. The number of MMA, Kme1, DMA, Kme2, Kme3, and mixed PSMs are shown. Mixed PSMs contained a mixture of methyl marks on the same peptide (e.g., MMA and DMA). The percolator q-value cutoff used to estimate the methyl FDR is also shown for each technique.

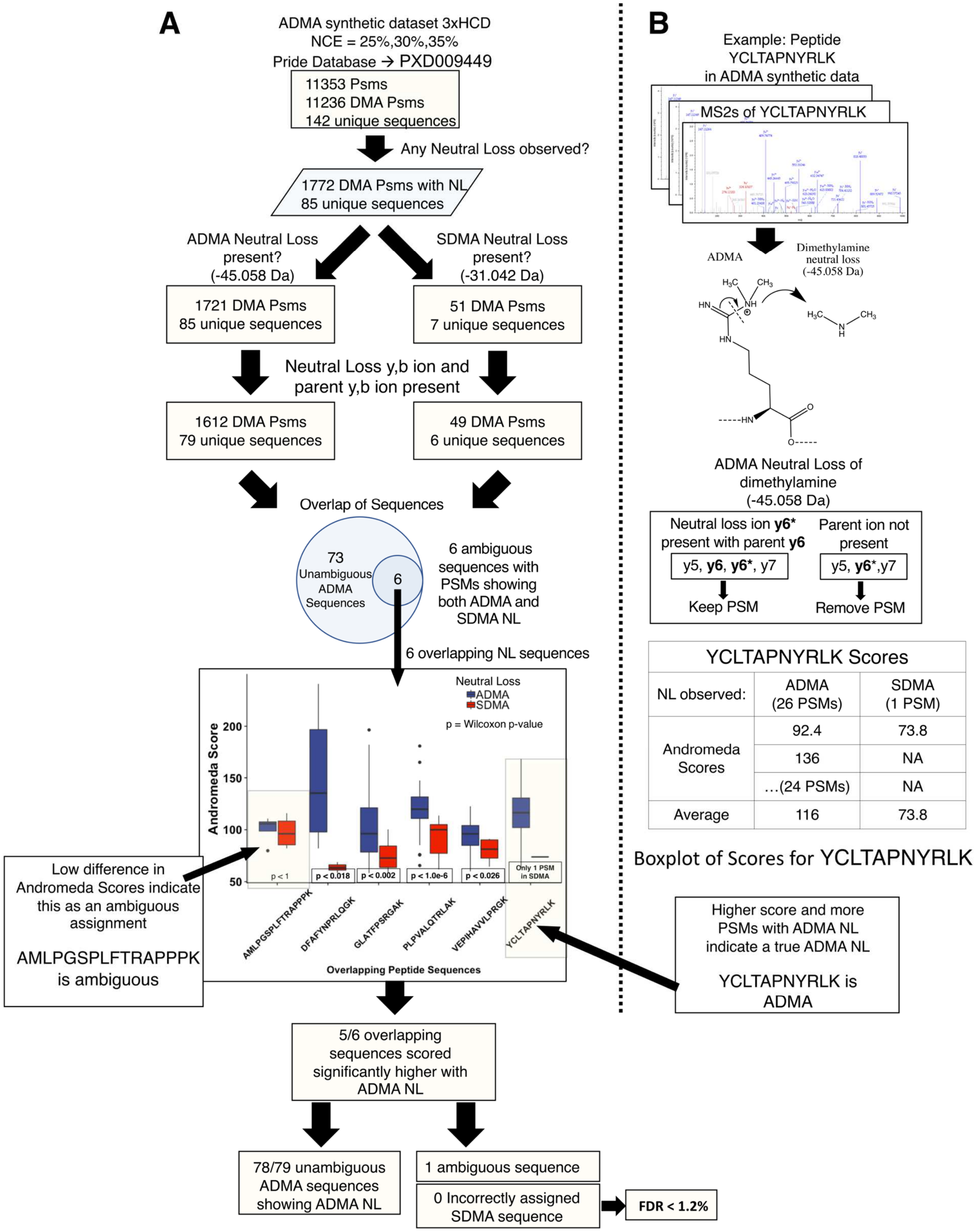
A
ADMA synthetic dataset 3xHCD
NCE = 25%,30%,35%
Pride Database → PXD009449
11353 Psms
11236 DMA Psms
142 unique sequences
Any Neutral Loss observed?
1772 DMA Psms with NL
85 unique sequences
ADMA Neutral Loss present? (-45.058 Da)
SDMA Neutral Loss present? (-31.042 Da)
1721 DMA Psms
85 unique sequences
51 DMA Psms
7 unique sequences
Neutral Loss y,b ion and parent y,b ion present
1612 DMA Psms
79 unique sequences
49 DMA Psms
6 unique sequences
Overlap of Sequences
73 Unambiguous ADMA Sequences
6
6 ambiguous sequences with PSMs showing both ADMA and SDMA NL
6 overlapping NL sequences
Neutral Loss
ADMA
SDMA
p = Wilcoxon p-value
Andromeda Score
200
150
100
50
p < 1
p < 0.018
p < 0.002
p < 1.0e-6
p < 0.026
Only 1 PSM in SDMA
AMLPGSPLFTRAPPPK
DFAFYNPRLQGK
GLATFPSRGAK
PLPVALQTRLAK
VEPIHAVVLPRGK
YCLTAPNYRLK
Overlapping Peptide Sequences
Low difference in Andromeda Scores indicate this as an ambiguous assignment
AMLPGSPLFTRAPPPK is ambiguous
5/6 overlapping sequences scored significantly higher with ADMA NL
78/79 unambiguous ADMA sequences showing ADMA NL
1 ambiguous sequence
0 Incorrectly assigned SDMA sequence
FDR < 1.2%
B
Example: Peptide YCLTAPNYRLK in ADMA synthetic data
MS2s of YCLTAPNYRLK
ADMA
Dimethylamine neutral loss (-45.058 Da)
ADMA Neutral Loss of dimethylamine (-45.058 Da)
Neutral loss ion y6* present with parent y6
y5, y6, y6*, y7
Keep PSM
Parent ion not present
y5, y6*,y7
Remove PSM
YCLTAPNYRLK Scores
NL observed:
ADMA (26 PSMs)
SDMA (1 PSM)
Andromeda Scores
92.4
73.8
136
NA
…(24 PSMs)
NA
Average
116
73.8
Boxplot of Scores for YCLTAPNYRLK
Higher score and more PSMs with ADMA NL indicate a true ADMA NL
YCLTAPNYRLK is ADMA

**Fig. S3. Andromeda identifies ADMA through neutral loss of dimethylamine**

A) Workflow diagram of the steps used to validate Andromeda searches for the neutral loss of dimethylamine in a synthetic ADMA dataset (3) (PXD009449). Roughly 140 synthetic peptides contained a single, non-terminal dimethyl ADMA. Starting from the Andromeda output msms.txt, peptides that exhibited neutral loss were annotated with either ADMA (dimethylamine) and/or SDMA neutral loss (methylamine) (4–6). We then required that each neutral loss ion be accompanied by the original b/y ion (e.g., y6* was kept if and only if y6 was also present). Our rationale for this filter was that neutral loss ions are typically weaker than the original b/y ion, so neutral loss ions not accompanied by the original b/y ion are likely to be false positive identifications. After this filtering, 73 peptides showed ADMA neutral loss and 6 peptides showed evidence of both ADMA and SDMA neutral losses. For the 6 ambiguous peptides, the mean Andromeda Score of the PSMs was calculated for all ADMA and SDMA PSMs. The boxplot in A) shows that for 6 of 7 synthetic methylated peptides their mean Andromeda Scores allowed clear assignment of ADMA. Only one ambiguous peptide which had similar Andromeda scores for ADMA and SDMA neutral loss PSMs could not be assigned. An identical process was used to process the synthetic SDMA dataset which also contained roughly 140 peptides. As a control, the unmodified synthetic dataset was also searched on Andromeda and no methylamine/dimethylamine neutral losses were detected (data not shown). For our own data, because MMA can also produce a neutral loss of methylamine, we imposed an additional restriction for mixed peptides containing both MMA and SDMA that that the SDMA site be localizable by the neutral loss ions. If the SDMA site could not be localized by neutral loss ions, the neutral loss was rejected. All reported neutral loss masses are charge +1.

B) Example calculation for one synthetic ADMA peptide, YCLTAPNYRLK. MS2 spectra of YCLTAPNYRLK were searched for ADMA and/or SDMA neutral losses. Spectra that contained neutral losses were subjected to the requirement that a neutral loss/original y ion pair must be present for the neutral loss to be considered. YCLTAPNYRLK had 26 PSMs with ADMA neutral losses and only 1 PSMs with SDMA neutral losses. The ADMA PSMs had an average Andromeda score of 116, whereas the SDMA PSM had a score of 73.8. Because ADMA PSMs were substantially more numerous (26 to 1) and the ADMA Andromeda scores were much higher than the SDMA Andromeda score suggested that YCLTAPNYRLK is ADMA modified. All reported neutral loss masses are charge +1.

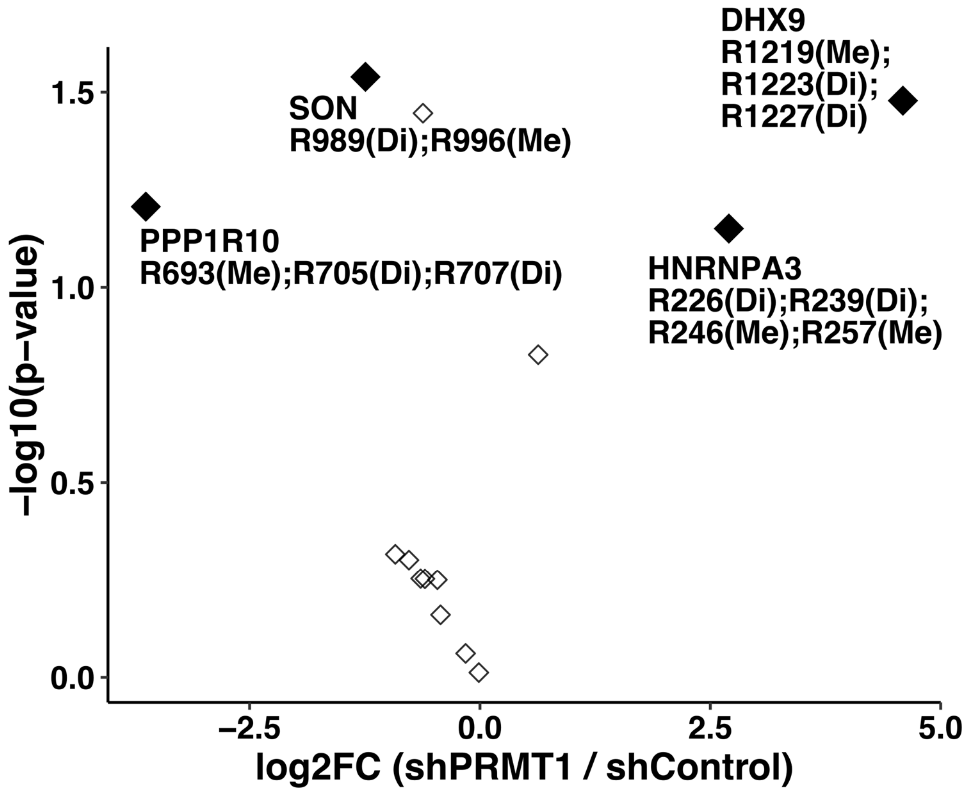


**Fig. S4. Quantification of mixed methyl arginine SCX peptides**

A) Volcano plot of quantified mixed methyl arginine peptides identified by SCX from 293T cells expressing shControl or shPRMT1. Mixed peptides contained both an MMA site and a DMA site on the same peptide. LFQ values were $log_2$ transformed, median normalized, and subjected to a permutation t-test in Perseus with $q < 0.05$, and $S_0 = 0.5$. Filled diamonds represent peptides that have a q-value < 0.05.

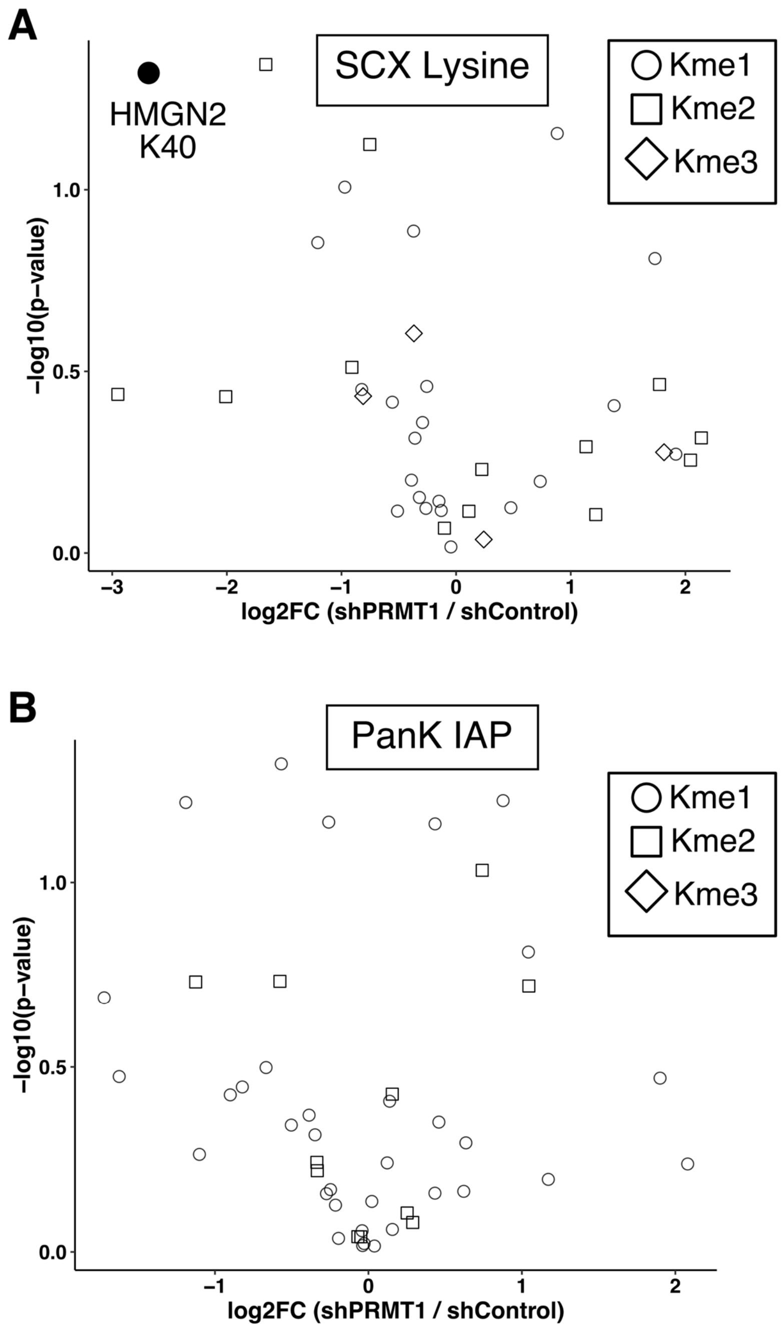


**Fig. S5. Lysine methylation is minimally affected by knockdown of PRMT1.**

A) Volcano plot of quantified methyl lysine peptides enriched by SCX on 293T cells expressing shControl and shPRMT1. LFQ values were $log_2$ transformed, median normalized, and subjected to a Perseus permutation-based t-test to asses significance with parameters $q < 0.05$ and $S_0$ =

0.5. Filled symbols represent a q-value < 0.05. Only one site, HMGN2 K40, was found to significantly change upon PRMT1 knockdown.

B) Similar to A) but for PanK IAP enriched methyl lysine peptides. No significant changes were observed for PanK IAP upon knockdown of PRMT1.

| Methyl Site | Peptide Sequence and Neutral Loss Assignment | DMA log2FC SCX | Q-value |
|---|---|---|---|
| G3BP1 R447; R460 | DMA SDMA<br>GPPRGGMVQKPGFGVGRGLAPR | -1.27 | 0.14 |
| G3BP1 R460 | SDMA<br>GGMVQKPGFGVGRGLAPR | 3.05 | 0.38 |
| FUS R216; R218 | SDMA DMA<br>GGRGRGGSGGGGGGGGGYNR | -1.10 | 0.18 |
| FUS R218 | DMA<br>GRGGSGGGGGGGGGYNR | 3.89 | 0.12 |
| CNBP R32 | DMA<br>SRGRGFQFVSSSLPDICYR | 1.26 | 0.37 |
| CNBP R32; R34 | DMA DMA There was an unlocalized SDMA neutral loss here<br>SRGRGFQFVSSSLPDICYR | -1.89 | 0.13 |
| CNBP R34 | DMA<br>GRGFQFVSSSLPDICYR | 0.84 | 0.37 |

**Fig. S6. Missed cleavage patterns of quantified methyl peptides demonstrate methylation changes upon PRMT1 knockdown.**

Quantified SCX peptides with missed cleavage variants from 293T cells expressing either shPRMT1 or shControl are displayed, along with $\log_2$ fold change of shPRMT1/shControl, permutation t-test q-values, and neutral losses if present. Examination of missed cleavage peptides improved confidence in methylation changes upon PRMT1 knockdown. (top) The missed cleavage peptide G3BP1 R447(DMA);R460(SDMA) was downregulated in PRMT1 knockdown cells, but the fully tryptic peptide G3BP1 R460(SDMA) was upregulated in PRMT1 knockdown cells. This suggests that R447 was demethylated in PRMT1 knockdown cells, leading to cleavage at R447 and therefore increased abundance of the G3BP1 R460(SDMA) peptide. We did not observe neutral losses to assign R447 as either ADMA or SDMA modified. (middle) The missed cleavage peptide FUS R216(SDMA);R218(DMA) was downregulated in PRMT1 knockdown cells, but the fully cleaved peptide FUS R218(DMA) was upregulated in PRMT1 knockdown cells. This suggests that FUS R216(SDMA) was demethylated in PRMT1 knockdown cells, leading to cleavage at R216 and therefore increased abundance of tryptic peptide FUS R218. (bottom) The missed cleavage peptide CNBP R32(DMA);R34(DMA) was downregulated in PRMT1 knockdown cells, but both the fully cleaved peptide CNBP R34(DMA) and the missed cleavage peptide CNBP R32(DMA) were upregulated in PRMT1 knockdown cells. This suggests that CNBP exists in doubly dimethylated form in shControl cells but in either of two singly dimethylated forms in shPRMT1 cells.
